# TEAD1 is a novel regulator of NRF2 and oxidative stress response in cardiomyocytes

**DOI:** 10.1101/2025.04.06.647438

**Authors:** Rajaganapathi Jagannathan, Jeongkyung Lee, Evan DeVallance, Vinny Negi, Varun Mandi, Hridyanshu Vyas, Feng Li, Patrick J. Pagano, Vijay Yechoor, Mousumi Moulik

## Abstract

**BACKGROUND:** TEAD1, the mammalian Hippo pathway-regulated transcription factor, plays a critical and non-redundant role in maintaining cardiomyocyte (CM) homeostasis. However, the specific cellular pathways regulated by TEAD1 in CMs remain poorly defined. We hypothesized that TEAD1 has an essential, cell-autonomous role in the CM oxidative stress response by directly regulating the transcription of NRF2, the master regulator of oxidative stress response.

**METHODS AND RESULTS:** Tamoxifen-induced conditional CM-specific TEAD1 deletion in adult mice leads to acute heart failure (HF) and altered expression of antioxidant genes. In silico analysis of publicly available RNA-seq data from human hearts with end-stage dilated (DCM) and ischemic (ICM) cardiomyopathy revealed significant downregulation of TEAD1 transcript levels and a positive correlation between TEAD1 and NRF2 gene expression. ChIP-seq and ATAC-seq in adult mouse hearts confirmed TEAD1 occupancy at promoter/enhancer elements within open chromatin regions of multiple antioxidant genes, including NRF2 and its targets. Ex vivo and in vitro TEAD1 knockout in primary neonatal and adult murine CMs, as well as in H9C2 cells, resulted in significantly increased cellular and mitochondrial ROS le, accompanied by a marked decrease in NRF2 expression and promoter-luciferase activity, under both basal and oxidative stress conditions. Mosaic, conditional deletion of TEAD1 in ∼40–50% of murine heart CMs provided a novel in vivo model for studying TEAD1-regulated pathways in the heart, independent of the confounding effects of HF. This model demonstrated reduced NRF2 expression and heightened oxidative stress in neonatal and adult TEAD1 mosaic knockout hearts. Notably, 8OHdG staining identified oxidative DNA damage in TEAD1-deficient CMs compared to TEAD1-expressing CMs within the mosaic knockout hearts. Upon in vivo AngII infusion, TEAD1 mosaic knockout hearts showed a significant increase in oxidative stress markers and an impaired NRF2 response. Overexpression of human TEAD1 restored NRF2 activity and mitigated ROS accumulation in TEAD1 knockout CMs in vitro. Furthermore, TEAD1 deletion in human iPSC-derived CMs resulted in increased oxidative stress and downregulation of NRF2 expression and functional activity, confirming the requirement of TEAD1 in NRF2-mediated oxidative stress response in human CMs. Collectively, these findings establish that TEAD1 is essential for NRF2 expression and activity under both basal and AngII-induced conditions and plays a crucial role in the oxidative stress response in CMs.

**CONCLUSIONS:** TEAD1 is a cell-autonomous, direct transcriptional regulator of NRF2 and the cardiomyocyte (CM) oxidative stress response. Its gene expression, which directly correlates with NRF2 transcript levels in the human myocardium, is significantly downregulated in human end-stage heart failure, potentially compromising the oxidative stress response in the failing heart.

## Introduction

Heart failure is a major cause of morbidity and mortality, accounting for over 13% of annual deaths worldwide^1^. While its etiologies are diverse—ranging from myocardial infarction and hypertension to cardiomyopathy, myocarditis, and valvular heart disease—disruptions in cardiomyocyte stress response pathways are a common feature underlying many of these conditions. Oxidative stress, endoplasmic reticulum (ER) stress, and hypoxic stress pathways are frequently activated in heart failure.

Substantial clinical and experimental evidence indicates that oxidative stress is a prevalent factor in heart failure, characterized by an accumulation of reactive oxygen species (ROS) resulting from excessive production and impaired antioxidant scavenging^2,3^. Oxidative stress leads to further impairment in cellular responses by protein and lipid modifications, DNA damage, and, eventually, cellular death^4,5^. The transcription factor NRF2 (*NFE2L2*) has been identified as a master regulator of the antioxidant response, though therapies aimed at augmenting NRF2 activity have not yet reached clinical application^6,7^. Advancing our understanding of the upstream regulators of NRF2 and the mechanisms underpinning a robust antioxidant response may uncover novel pathways to enhance cardiomyocyte resilience, offering new opportunities to prevent and mitigate heart failure.

The mammalian hippo signaling pathway is an evolutionarily conserved signaling pathway that regulates organ size, cellular growth, proliferation, apoptosis, and maintenance of mature function^8^. It comprises a core kinase cascade that phosphorylates and inhibits YAP and TAZ, coactivators for the TEAD family of transcription factors. The hippo-TEAD1 pathway plays a significant role in cardiac development and function^9^. Hippo upstream regulators have also been shown to be involved in cardiac remodeling^9–11^ and regeneration after myocardial infarction^12^.

TEAD1 is a critical transcription factor essential for heart development and function across all stages, from early embryogenesis to perinatal period and postnatal adulthood. Deletion of TEAD1 during embryonic development is lethal due to impaired cardiogenesis^13^. Similarly, perinatal deletion of TEAD1 results in reduced cardiomyocyte proliferation and fatal neonatal cardiomyopathy. Inducible conditional deletion of TEAD1in adult CMs leads to rapid onset acute heart failure and mortality within a few weeks, accompanied by impaired excitation-contraction coupling and reduced TEAD1 regulated SERCA2A activity^14^. Furthermore, TEAD1 is required for normal mitochondrial function in cardiomyocytes^15,16^. Diminished TEAD1 transcriptional activity, resulting from perinuclear trapping, has been implicated as a key mechanism in a knock-in mouse model of human DCM caused by a Lamin A/C mutation. However, the specific cellular pathways regulated by TEAD1 in CMs remain poorly defined. We hypothesized that TEAD1, a master regulator of mitochondrial homeostasis and mature cardiomyocyte function, also regulates stress pathways, including the oxidative stress pathway, to maintain cardiomyocyte resilience. The acute and severe HF observed following both constitutive MYH6-Cre and tamoxifen-inducible MYH6-MerCreMer-mediated CM-specific conditional deletion of TEAD1 precluded the delineation of primary TEAD1-regulated functional pathways, independent of the confounding effects of HF-induced secondary remodeling. To overcome this hurdle and test our hypothesis, we developed and characterized a mosaic TEAD1 deletion mouse model. In this model, TEAD1 is deleted in approximately 40–50% of CMs, allowing the mice to maintain normal ejection fraction and avoid HF during the study period. This approach enabled detailed molecular analysis of TEAD1-regulated pathways without the interference of heart failure-related secondary changes.

In this study, we demonstrate a significant downregulation of TEAD1 gene expression in the hearts of patients with end stage heart failure due to non-ischemic dilated cardiomyopathy (DCM) and ischemic cardiomyopathy (ICM) and a strong positive correlation between TEAD1 transcript levels and the expression of multiple antioxidant genes, including NRF2, in human hearts. In vitro and in vivo experiments demonstrate that TEAD1 is a direct transcriptional regulator of NRF2 in cardiomyocytes, and its deletion leads to a decrease in NRF2 transcript, and protein levels, and ARE promoter activity, accompanied by an increase in ROS and oxidative stress. In vivo, mosaic deletion of TEAD1 in ∼40-50% of cardiomyocytes reveal that only the TEAD1-deleted cardiomyocytes experience oxidative stress at baseline, and this is significantly worsened when stressed with angiotensin II. TEAD1 deletion in h-iPSC-derived CMs confirms requirement of TEAD1 for NRF2 expression and function. Collectively, these results establish the novel finding that TEAD1 regulates NRF2 and the antioxidant response in CMs, playing a cell-autonomous role in preventing oxidative stress and maintaining cardiomyocyte resilience. Targeting this pathway may provide novel therapeutic opportunities to mitigate cardiac damage and improve heart function in heart failure.

## Result

### TEAD1 expression in human hearts is significantly correlated with antioxidant gene expression in heart failure

We have previously shown that TEAD1 is required for normal cardiac function, and in vivo deletion of TEAD1 in mouse cardiomyocytes leads to lethal heart failure^13–15^. In addition, our data on TEAD1 protein expression from a small cohort of DCM patients revealed downregulation of TEAD1, suggesting possible TEAD1 alteration in human HF^14^. However, TEAD1 expression in large cohorts of human heart failure has not been previously reported. Hence, to determine if the expression of TEAD1 and Hippo-TEAD pathway genes is altered across the spectrum of human heart failure, we analyzed their expression in publicly available RNA-seq data sets (GSE141910^17,18^ and GSE116250^19,20^) from patients with end-stage heart failure. TEAD1 expression was significantly decreased across 216 DCM heart failure samples compared to control hearts with normal function, along with significant alteration of many Hippo Pathway (KEGG pathway) genes (Fig.1a-b). Analysis of differentially expressed genes (DEGs) that were common in both data sets (Fig. 1c) using Ingenuity Pathway Analysis (IPA) revealed that nuclear factor erythroid 2-related factor2 (NRF2)-mediated oxidative stress response and glutathione redox reaction pathways were significantly downregulated while the inhibitory Hippo pathway signaling was upregulated (Fig.1d). Similar downregulation of TEAD1 and the NRF2-mediated oxidative stress response and glutathione redox reaction pathways were also noted in ICM samples (GSE116250) as compared to their controls (Supplementary Fig. S1a-b). Our prior gene expression data from TEAD1 knock-out mouse hearts had revealed differential enrichment of oxidative stress response pathway genes, suggesting possible TEAD1 regulation of cardiac oxidative stress response, though it could not be distinguished from the confounding effect of a secondary HF molecular signature^15^. To determine if there is a correlation between TEAD1 and oxidative stress response gene expression in human hearts, we performed Pearson’s linear correlation analyses between transcript levels of TEAD1 and oxidative stress response genes, which revealed a significant, positive correlation between TEAD1 and several key NRF2-mediated oxidative stress response pathway genes, including moderate correlation with NRF2 (NFE2L2) (R=0.4, p<0.0001), HMOX1 (R=0.6, p<0.0001), NQO1 (R=0.6, p<0.0001), SOD1 (R=0.5, p<0.0001) and SOD2 (R=0.5, p<0.0001) and a weak correlation with SLC7A11 (R=0.1, p=0.02) across all heart samples in these human data sets (Fig. 1e). This correlation between TEAD1 and NRF2 and its target genes, suggested that TEAD1 could be a regulator of NRF2-mediated oxidative stress response pathway in the heart and that this may underpin a critical function of TEAD1 in cardiomyocytes.

**Figure 1:**
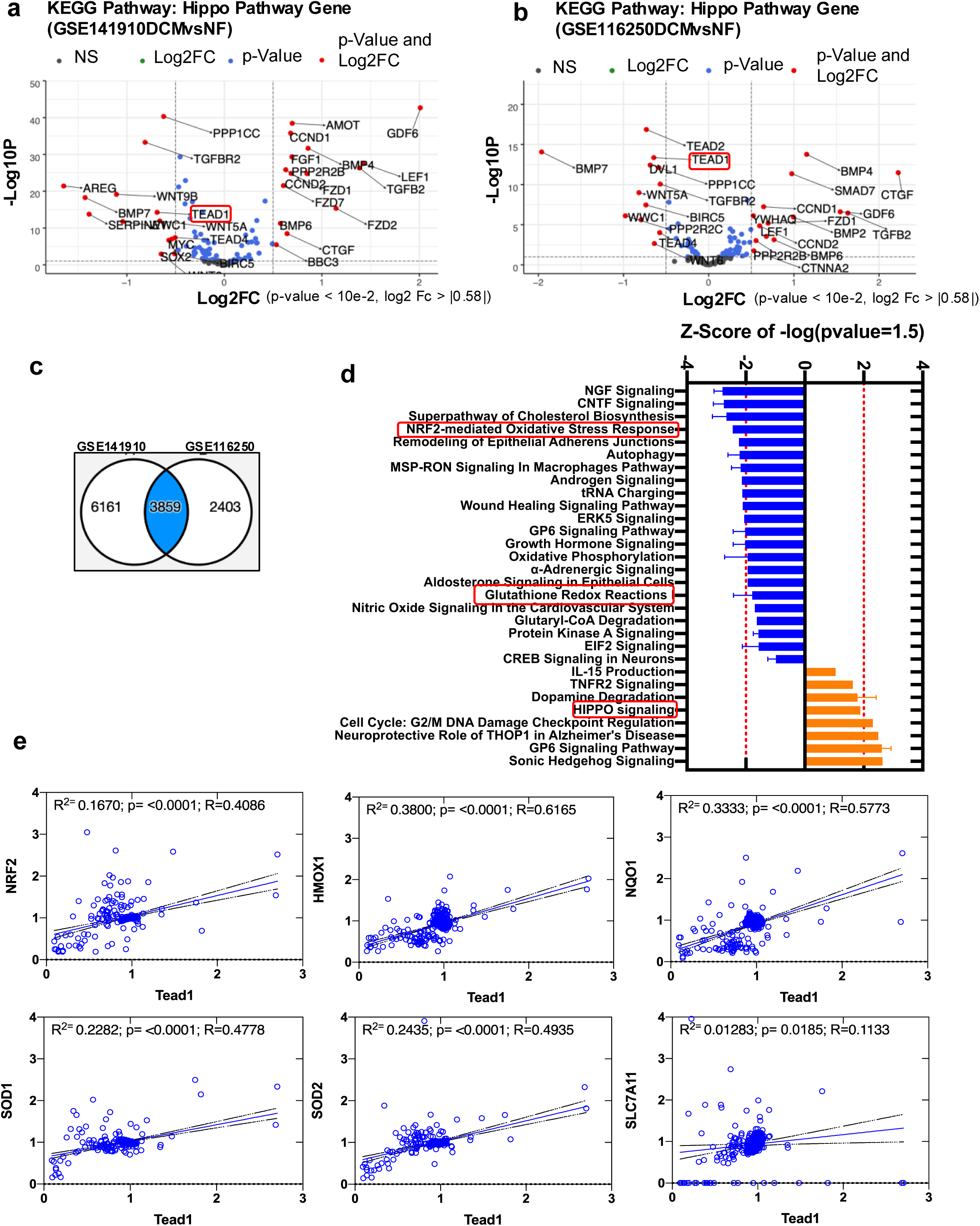
TEAD1 expression is positively correlated with antioxidant gene expression in human heart failure. RNA-seq datasets from human heart samples from GSE141910 and GSE116250 were reanalyzed, normalized, and fold-expression compared to their respective non-failing heart controls was calculated. **(a-b)** Volcano plots showing the Hippo signaling pathway (KEGG) significantly regulated in (**a**) GSE141910 - dilated cardiomyopathy (DCM) hearts (n=166) compared to non-failing (NF) hearts (n=166) and (**b**) GSE116250 - DCM hearts (n=37) compared to NF hearts (n=14). Significantly up/down-regulated genes are represented by red dots, significant, but log2FC < 0.58 - up/down-regulated genes by blue dots, and no significant changes by gray dots (p-value < 10e-2, log2 FC > |0.58|). **c)** Venn diagram showing the differentially expressed genes (DEGs) (in blue) significantly regulated (FDR adjusted p-value < 0.05) in DCM compared to NF hearts in both in both data sets (GSE141910 and GSE116250). **d)** Bar graph representing Ingenuity Pathway Analysis (IPA) of significantly inhibited (blue bars) or activated (orange bars) signaling pathways from DEGs common to GSE141910 and GSE116250 datasets. The red dotted line indicates a z-score cutoff of 2. **e)** NRF2 pathway gene expression (including NFE2L2 (NRF2), HMOX1, NQO1, SOD1, SOD2, and SLC7A11) in all heart samples across the datasets (GSE141910 and GSE116250) (n=397 includes DCM and NF controls). Correlation analyses with TEAD1 expression are shown, with Pearson’s correlation coefficient (r) and p-values provided for each analysis.

### TEAD1 directly regulates genes encoding antioxidant proteins and the NRF2 signaling pathway

To unravel regulatory mechanisms underlying this novel correlation between TEAD1 and NRF2/oxidative stress response gene transcripts, and to determine if TEAD1 was a direct transcriptional regulator of NRF2 in the adult heart, we performed chromatin immunoprecipitation using anti-TEAD1 antibody followed by sequencing (ChIP-seq) in 10-week-old wild-type (WT) mouse hearts and analyzed the genome-wide occupancy of TEAD1. TEAD1 binding was identified in 28,625 peaks across the genome. Mapping TEAD1-bound peaks from -5 kb to +5 kb from the transcription start site (TSS) of genes showed that TEAD1 binding is predominantly localized within 2kb around the TSS (Fig. 2a and Supplementary Fig. S2a). Most peaks were detected in the proximal promoter (<1kb – 27.23%) or in the distal intergenic regions 25.01%) and 13.36% distributed evenly over the 1-5 kb upstream promoter region, and 10.5% in the 1^st^ intron (Fig. 2b). Other binding regions included 1.37%, 1.82%, and 19.57% of the peaks within 1st Exon, other Exon, and other intronic regions respectively (Fig. 2b). Less than 1% binding was found in the 5’ UTR (0.12%), 3’UTR (0.98%), and downstream (<=300 kb) (0.05%). The peaks mapped were analyzed using MEME de novo transcriptional factor (TF) suite to identify enriched TF motifs, which revealed that, as expected, TEAD (1, 2, and 4) binding motif was the most significantly enriched, lending confidence to the quality of the ChIP-seq. Motifs for HAND1, MEFs (2A, 2B, and 2C), NFI (A, B, and C), all with known critical roles in cardiomyocyte development and function were the next 3 enriched motifs (Fig. 2c), suggesting a likely combinatorial role between TEAD1 and these TFs in transcriptional regulation. Using the Genomic Regions Enrichment of Annotations Tool (GREAT), we annotated these peaks with their closest genes and then performed a functional enrichment analysis (Supplementary Table 1). The 28,625 peaks were annotated to 13,377 protein-coding region genes (Supplementary Table 2). These peaks mapped to genes enriched for pathways related to multiple signaling pathways including cardiac hypertrophy signaling, NFAT related cardiac hypertrophy, G-protein coupled receptor signing and NRF2-mediated oxidative stress response signaling (Fig. 2d). Gene ontology (GO) biologic function enrichment analysis for cellular processes showed significant enrichment of stress response, including response to reactive oxygen species (H2O2), response to oxidative stress, protein folding, and cellular response to DNA damage stimulus (Supplementary Fig. S2c-d).

**Figure 2:**
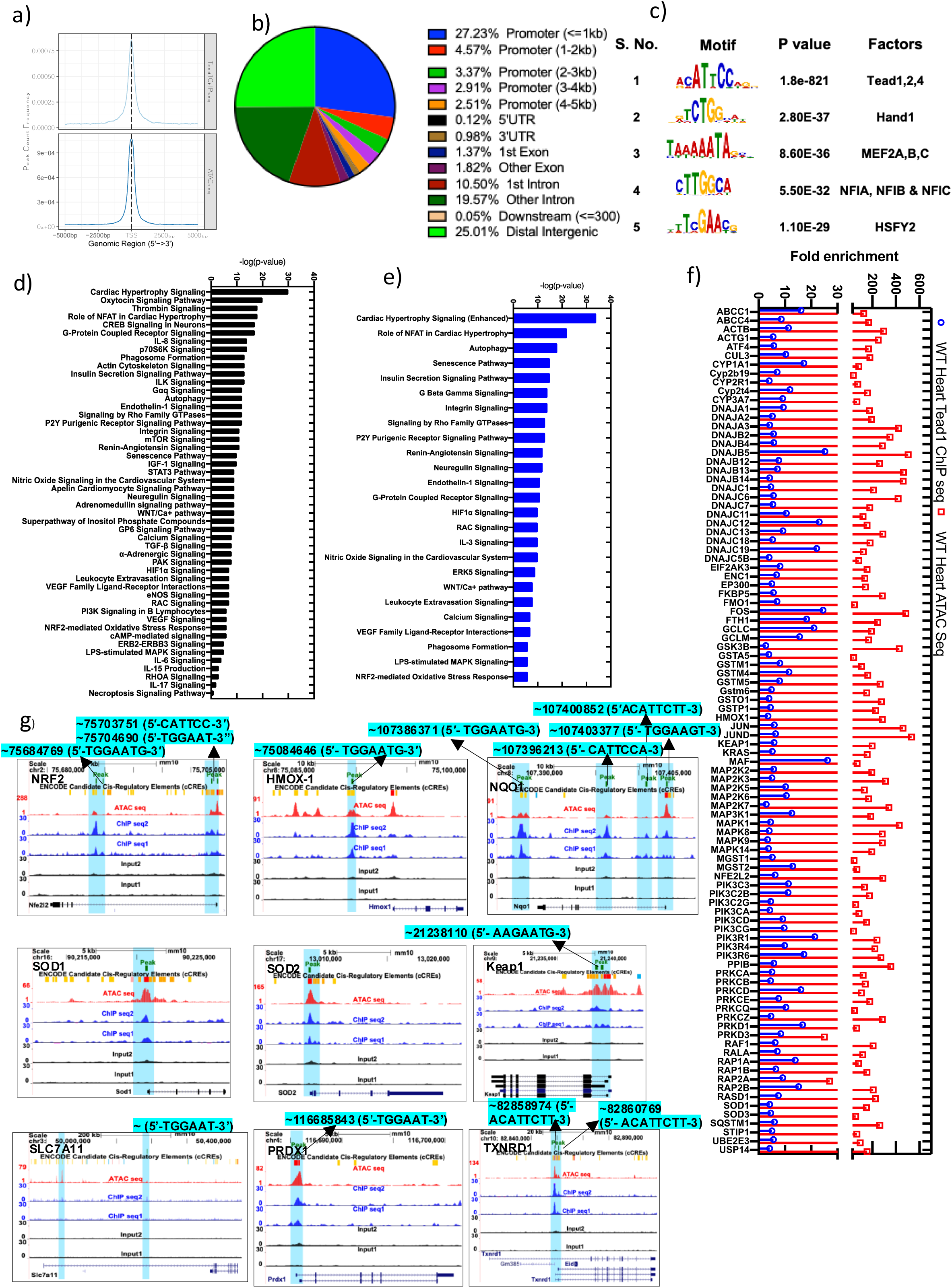
Tead1 is a direct transcriptional activator of antioxidant genes in murine heart. **a)** Composite plots of normalized Tead1 ChIP-seq and ATAC-seq signals in 10-week-old WT hearts, showing 5 kb centered around transcription start site (TSS). **b)** Genomic distribution of Tead1 ChIP binding in 10-week-old WT hearts. **c)** Motif analysis for enriched Tead1 ChIP-seq peaks. **d)** Top 50 significantly enriched pathways binding in the Tead1 ChIP-seq, with bar heights indicating the significance of binding. **e)** Top 25 significant common pathways between Tead1 ChIP-seq and ATAC-seq. **f)** Combined enriched Tead1 ChIP-seq and open ATAC-seq peak values for NRF2 pathway genes. **g)** Enriched Tead1 ChIP-seq and open chromatin (ATAC-seq) tracks for the genomic regions of NRF2, HMOX1, NQO1, KEAP1, SOD1, SOD2, PRDX1, and TXNRD1, showing Tead1 binding motifs in their peak regions.

To identify TEAD1 binding in transcriptionally active regions of the chromatin, we performed Assay for Transposase-Accessible Chromatin with high-throughput sequencing (ATAC seq) on WT 10 weeks old mice heart tissue to analyze genome-wide open chromatin region in adult hearts. We found 52,997 transposase accessible peaks suggesting regions of open chromatin and active transcription. As seen with the TEAD1 ChIP-seq, open chromatin regions were localized predominantly within 2kb around the TSS (Fig. 2a and Supplementary Fig. S2a), with promotor <=1kb 28.41%, 11.87% evenly distributed over the 1-5 kb upstream promoter promotor, and 10.08% in the 1^st^ intron. 1.19%, 1.27%, 2.05%, 20.08 and 24.81% of the peaks binding within 3’UTR, 1st Exon, other Exon, and other intron distal intergenic regions, respectively (Supplementary. Fig 2b). Less than 1% percentages of binding were found in the 5’UTR (0.12%) and downstream (<=300) (0.09%). Ingenuity pathway analysis of ATAC-seq peaks revealed open chromatin was enriched for genes associated with pathways in protein ubiquitination, senescence, sirtuin signaling, EIF, AMPK signing pathways, and relevant to this study, NRF2-mediated oxidative stress response signaling was also enriched (Supplementary Fig. S2e). By cross-comparing TEAD1 ChIP-seq targets and ATAC-seq open chromatin genes, several common pathways were identified that would represent likely functional direct transcriptional targets of TEAD1 in adult heart. The common pathways include cardiac hypertrophy signaling, NFAT cardiac hypertrophy, autophagy, GPCR signaling, and NRF2-mediated oxidative stress-responsive signaling, among others (Fig. 2e). In NRF2-mediated oxidative stress-responsive signaling, 51 genes were commonly enriched between TEAD1 ChIP-seq and ATAC-seq (Fig. 2f). Overlapping TEAD1 binding and ATAC-seq peaks (transcriptionally active open chromatin regions) were noted in the promoters of critical antioxidant genes, including NRF2, HMOX1, NQO1, SOD1&2, KEAP, SLC7A11, PRDX1, and TXNRD1 (Fig. 2g). with these regions containing TEAD1 binding motifs. Collectively, these data strongly indicate that TEAD1 is a direct transcriptional regulator of the antioxidant response in the heart.

### Cell-autonomous TEAD1 function is required for normal antioxidant function in H9C2 cardiomyocyte cell line

In order to test the cell-autonomous requirement of TEAD1 in CM oxidative stress response, under in vitro conditions., we generated a stable CRISPR-Cas9-mediated *Tead1* KO in H9C2, the rat ventricular cardiomyocyte cell line (Fig. 3a) Tead1-depletion led to a significant accumulation of reactive oxygen species, a 3.5 fold increase in cellular hydrogen peroxide (boronate assay), and a ∼40% increase in mitochondrial superoxide (mitosox assay) levels, as compared to control H9C2 cells (Fig. 3b-c). This demonstrated that the cell-autonomous function of TEAD1 was required to prevent ROS accumulation in the basal state. We next asked if TEAD1 is also required to mitigate ROS accumulation with disease-state-associated oxidative stress. To test this, we exposed *Tead1*KO and control H9C2 cells to several oxidative stress inducers. Exposure to 10uM DMNQ (ROS inducer), 10 µM antimycin A (complex III inhibitor), and 200µM H2O2 (oxidative stress-inducer) led to an expected increase in mitochondrial superoxide level (Mitosox assay) in the control cells compared to basal state. TEAD1 KO cells showed a significant increase of ∼30-70% (p<0.001 in mitochondrial superoxide with DMNQ and H2O2 as compared to controls (Fig. 3d-f), confirming TEAD1 requirement for manifesting robust oxidative stress response under conditions of oxidative stress.

**Figure 3:**
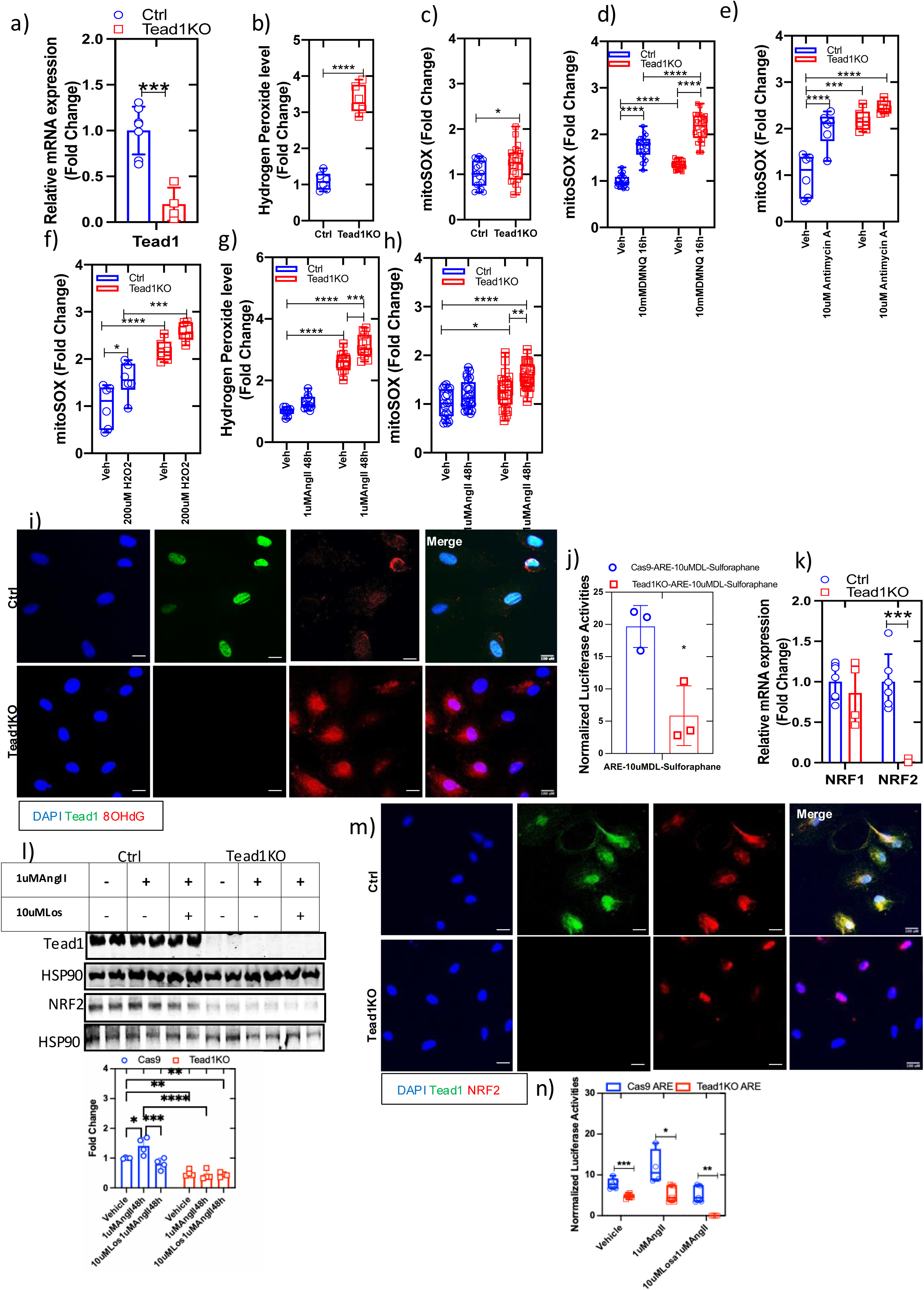
TEAD1 deletion in H9C2 cardiomyocyte cell line leads to a loss of NRF2 function and increased ROS. **a)** Expression of Tead1 gene expression by qPCR in Tead1 knockout (KO) stable H9C2 (TEAD1KO) and Cas9 scramble control sgRNA (Cas9) cell lines created using CRISPR-Cas9. **b)** Quantification of total ROS (Hydrogen Peroxide) levels measured using the coumarin boronate acid (CBA) assay in TEAD1KO and control H9C2 cells. **c)** Quantification of mitochondrial superoxide levels measured using the mitoSOX assay in TEAD1KO and control H9C2 cells. **d)** Quantification of mitochondrial superoxide levels measured using the mitoSOX assay in TEAD1KO and control H9C2 cells treated with vehicle or ROS inducer 10 mM DMNQ for 16 hours. **e)** Quantification of mitochondrial superoxide levels measured using the mitoSOX assay in TEAD1KO and control H9C2 cells treated 10 µM Antimycin A (mitochondrial complex III inhibitor) for 2 hours or vehicle (DMSO 0.01%). **f)** Quantification of mitochondrial superoxide levels measured using the mitoSOX assay in TEAD1KO and control H9C2 cells treated with vehicle or 200 µM H2O2 for 2 hours. **g)** Quantification of total ROS (Hydrogen Peroxide) levels measured using the CBA assay in TEAD1KO and control H9C2 cells challenged vehicle or 1 µM AngII for 48 hours. **h)** Quantification of mitochondrial superoxide levels measured using the mitoSOX assay in TEAD1KO and control H9C2 cells treated with 1 µM AngII for 48 hours or vehicle (PBS). **i)** Representative immunofluorescent images of 8-OHdG labeling in control (top) and TEAD1KO (bottom) H9C2 cells [anti-Tead1 (green), anti-8-hydroxydeoxyguanosine (anti-8-OHdG) (red), and nucleus-DAPI (blue) staining]. **j)** ARE-luciferase reporter assay to assess NRF2 activity in TEAD1KO and control H9C2 cell lines exposed to DL-Sulforaphane (SF) 10 μM for 24 hours. **k)** mRNA expression of NRF1 and NRF2 in control and TEAD1KO H9C2cells (n=4-6). **l)** Western blot analysis of TEAD1 and NRF2 protein levels in TEAD1KO and control H9C2 cells treated with vehicle, Angiotensin II (1 µM AngII) with or without antagonist (10 mM Losartan) for 48 hours, with quantification shown below. HSP90 was the loading control. **m)** Representative immunofluorescent images of NRF2 and TEAD1 in control (top) and TEAD1KO (bottom) H9C2 cells [anti-Tead1 (green), anti-NRF2 (red), and nucleus-DAPI (blue) staining]. **n)** ARE-luciferase reporter assay to assess NRF2 activity in TEAD1KO and control H9C2 cell lines treated with vehicle (PBS) or 200 nM AngII with or without antagonist (10 mM Losartan) for 48 hours (n=5-6). Statistical significance is indicated as follows: ***p<0.0005; **p<0.005; *p<0.05; no symbol indicates no significance, compared with control (Student’s t-test and Two-way ANOVA). Scale bar: 100 µm.

Cardiac remodeling is associated with oxidative stress^21,22^ and angiotensin II (AngII), a frequent mediator of remodeling, is a known inducer of pathologic oxidative stress^23^. When challenged with 1uM AngII for 48 hours, *TEAD1*KO cells displayed a significant increase of hydrogen peroxide and mitochondrial superoxide (∼100% and 45% respectively) (Fig. 3g-h). Additionally, ROS-induced DNA damage, as measured by 8-hydroxy-2′-deoxyguanosine (8-OHdG) immunostaining, was evident upon TEAD1 depletion indicating functionally significant oxidative stress (Fig. 3i). This suggested that TEAD1 activity is required to mitigate basal and induced oxidative stress in cardiomyocytes.

We next assessed NRF2 transcriptional activation function (ARE promoter-luciferase activity) and expression levels in TEAD1 KO cells, to determine if TEAD1 regulation of oxidative stress response was mediated through NRF2. NRF2 is the master antioxidant transcription factor that upregulates many genes encoding antioxidant enzymes and proteins by binding to the Antioxidant Response Element (ARE) motif in their promoters. In the basal state, NRF2 is inactive, bound to KEAP1 in the cytoplasm, and rapidly degraded. Inhibition of this interaction with sulforaphane releases NRF2, allowing nuclear translocation and subsequent activation of target gene transcription. Sulforaphane induced a 20-fold increase in NRF2-ARE-luciferase (ARE-Luc) activity in the control H9C2 cells but only about a 5-fold increase in the absence of TEAD1 (Fig. 3j). The transcript and protein for NRF2 were significantly decreased by over 90% and 50% respectively in *TEAD1*KO cells as compared to controls (Fig. 3k-l). On AngII challenge, control cells displayed an increase in NRF2 protein level in response to the oxidative stress induced by AngII, while *TEAD1*KO cells did not have any significant increase over baseline. The specificity of this effect was confirmed by its sensitivity to Losartan, the Angiotensin receptor 1 (ATR1) inhibitor in control cells (Fig. 3l). This was accompanied by a decrease in nuclear accumulation of NRF2 in *TEAD1*KO cells (Fig. 3m) and a lack of induction of ARE-luc activity upon exposure to AngII (Fig. 3n). This lack of an appropriate increase in NRF2 with AngII challenge would be expected to increase ROS accumulation and hence could be the mechanistic basis for the observed increase in ROS in *TEAD1*KO cells when exposed to AngII (Fig. 3g-h). These data demonstrated that Tead1 is required for NRF2-dependent antioxidant response to mitigate cellular ROS accumulation in both basal and stressed cardiomyocytes.

### Ablation of *TEAD1* impairs antioxidant regulation in primary neonatal cardiomyocytes

Previously published data from our lab and other reports^15,16^ demonstrated the non-redundant requirement of TEAD1 in CM function and specifically in maintaining normal mitochondrial homeostasis in vivo. Our global transcriptome analysis from TEAD1-deficient hearts revealed significant enrichment in oxidative stress/mitochondrial function genes^15^. Since some of these transcriptomic changes could be secondary to heart failure seen in TEAD1-deficient hearts, to determine the primary effect of TEAD1 deficiency on the global CM transcriptome, we performed inducible ex vivo TEAD1 deletion in neonatal CMs (iNCMs). We isolated neonatal cardiomyocytes from P1-2 day old *Tead1^F/F^* pups and infected them with Ad-GFP (control) or with Adenovirus carrying Cre-GFP (Ad-Cre-GFP) to delete *Tead1* (Fig. 4a and Supplementary Fig. S3a). This led to a >70% significant decrease in *Tead1* transcript in Ad-Cre-GFP infected *Tead1^F/F^* iNCMs, along with a significant decrease in transcripts for many cardiomyocyte markers, such as ANP, BNP, MEF2c and αActinin (Fig. 4b). Analysis of the global transcriptome of these TEAD1-deleted iNCMs revealed significant changes in gene expression with 1609 upregulated and 2555 downregulated transcripts (FDR adj. p value<=0.05; Fold change 1.2) as compared to Ad-GFP treated controls (Fig. 4c). An IPA analysis of these differentially expressed genes revealed NRF2-mediated oxidative stress response pathways to be enriched with loss of TEAD1 function, along with stress pathways, including p53 signaling, death receptor signaling, unfolded protein response, HIF1α signaling among others (Fig. 4d).

**Figure 4:**
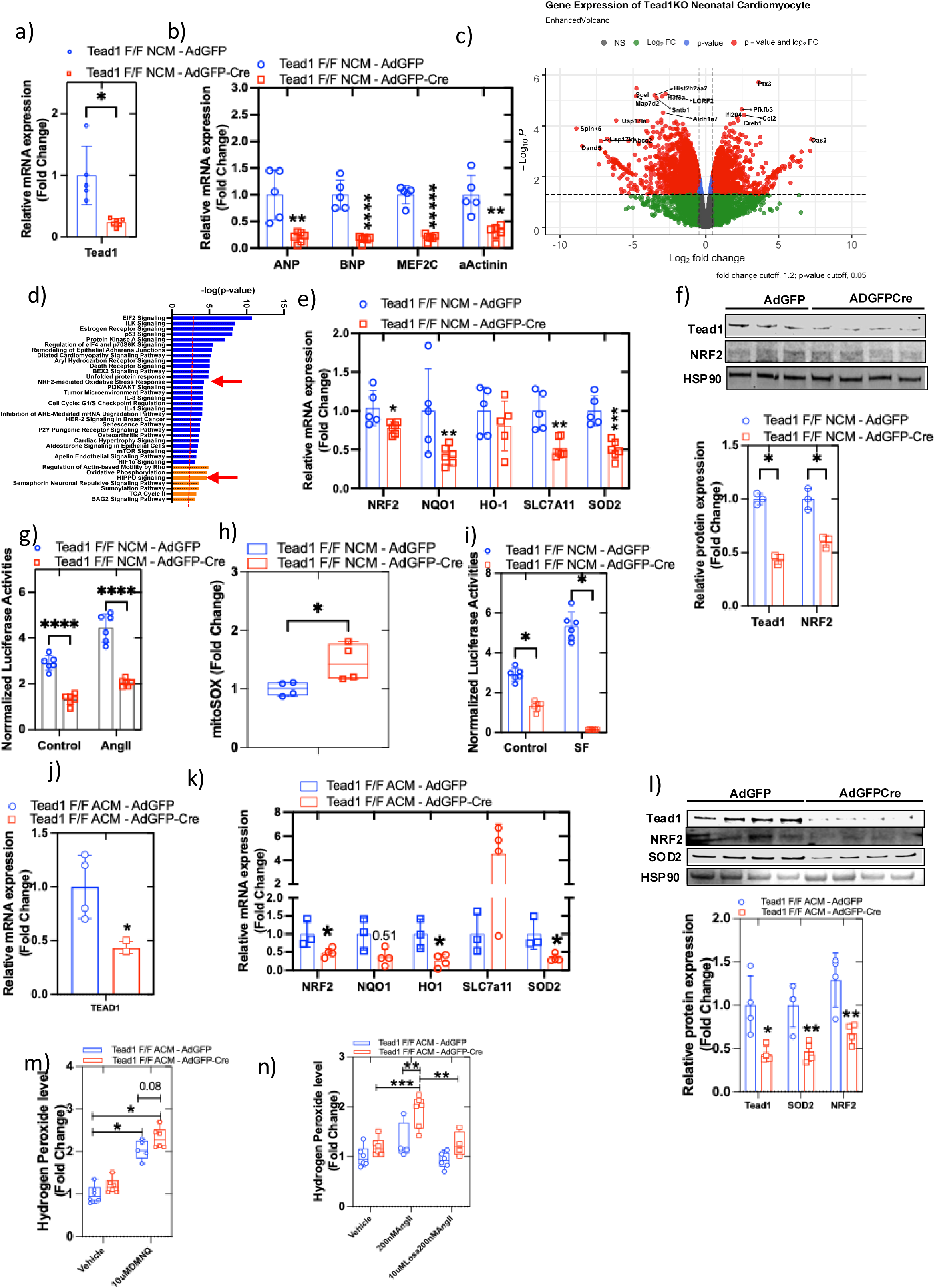
Cell autonomous function of TEAD1 is required in primary cardiomyocytes to prevent a loss of NRF2 function and increased ROS accumulation. **a-b)** Expression of TEAD1 (a) and other CM markers (b) by qPCR in isolated murine neonatal (P1) cardiomyocytes (iNCM) from Tead11f/f mice treated with adenoviral-GFP (AdGFP) [Control] and adenoviral-GFP-Cre (AdGFP-Cre) [TEAD1KO] and assessed after 72 hours. (n=5-6). **c)** RNA-Seq analysis of the global transcriptome of the TEAD1KO and Control iNCMs are shown in a volcano plot representation, with some of the Hippo signaling pathway genes highlighted. differentially expressed genes (DEGs) are represented by red dots (p-value < 0.05, log2 log2FC > |0.58|). **d)** Bar graph representing Ingenuity Pathway Analysis (IPA) of significantly activated (orange bars) and inhibited (blue bars) signaling pathways in TEAD1KO iNCMs, using DEGs. The red dotted line indicates a z-score cutoff of 2. **e)** Gene expression analysis of NRF2 and its target genes (NQO1, HO-1, SLC1A11, and SOD2) differentially expressed in TEAD1KO compared to control in iNCMs (n=5-6). **f)** Immunoblotting analysis of TEAD1 and NRF2 in control and TEAD1KO iNCMs, with quantification shown below. HSP90 was the loading control. **g)** ARE-luciferase reporter assay to assess NRF2 activity in TEAD1KO and control iNCMs exposed to vehicle (PBS) or to 200 nM AngII for 48 hours. **h)** Superoxide levels (mitoSOX assay) in TEAD1KO and control iNCMs. **i)** ARE-luciferase reporter assay to assess NRF2 activity in TEAD1KO and control iNCMs, treated with vehicle (DMSO 0.1%) or with Sulforaphane (SF) 10 μM for 24 hours. **j-k)** Expression of TEAD1 (j) and NRF2 and its target genes (NQO1, HO-1, SLC1A11, and SOD2) (k) by qPCR in isolated murine adult cardiomyocytes (iACM) from Tead11f/f mice treated with adenoviral-GFP (AdGFP) [Control] and adenoviral-GFP-Cre (AdGFP-Cre) [TEAD1KO] and assessed after 72 hours. (n=5-6). **l)** Western blotting analysis of control and TEAD1KO in iACM, for TEAD1, NRF2, and SOD2 protein levels, with quantification shown below. HSP90 was the loading control. **m)** Quantification of total ROS (hydrogen peroxide) in control and TEAD1KO iACMs challenged with the ROS inducer 10 µM 2,3-dimethoxy-1,4-naphthalenedione (DMNQ) for 2 hours. **n)** Quantification of total ROS (hydrogen peroxide) in control and TEAD1KO iACMs treated with Angiotensin II (1 µM AngII) or Ang II receptor antagonist (10 mM Losartan) for 48 hours. Values indicate fold changes in each group compared to the control group (±SEM). Statistical significance is indicated as follows: ***p<0.0005; **p<0.005; *p<0.05; no symbol indicates non-significant (n = 3–6 per group).

qRT-PCR and western blotting revealed that NRF2 transcript and protein were significantly decreased in TEAD1-deficient NCMs compared to controls (Fig. 4e-f). This decrease in NRF2 protein was also accompanied by a decrease in the transcripts for NRF2 targets, such as NQO1, SOD2, and SLC7A11 in TEAD1-deficient cells (Fig. 4e). HO-1, another target of NRF2, had a trend to be lower in expression with TEAD1-deletion as compared to control, but did not reach statistical significance (Fig. 4e). This was consistent with a decrease in the activity of NRF2 bound to the anti-oxidative stress response element (ARE), as assessed by ARE-luc activity, in TEAD1-deficient iNCMs (Fig. 4g). Despite stimulation for 48 hours with 200M of angiotensin II, a known inducer of oxidative stress in cardiomyocytes^24^, the expected increase in ARE-luc activity, as observed in the controls, was completely abrogated in TEAD1-deficient iNCMs (Fig. 4g). This decreased NRF2 activity translated to a significant increase by ∼40% in ROS (mitochondrial superoxide) accumulation, quantitated by Mitosox assay, in TEAD1-deficient iNCMs (Fig. 4h). Sulforaphane induced an increase in NRF2-ARE-luc activity in the control NCMs but was not able to induce NRF2-ARE-luc activity in the absence of TEAD1 (Fig. 4i). These results demonstrate that cell-autonomous TEAD1 function is required for regular NRF2 expression, and antioxidant response in neonatal CMs under basal and stressed conditions.

### TEAD1 is required for NRF2 expression and mitigating oxidative stress in primary adult cardiomyocytes

To assess if cell-autonomous TEAD1 function is also required for NRF2-mediated antioxidant response in adult CMs, we generated inducible deletion of TEAD1 in adult CMs. We isolated primary cardiomyocytes from *Tead1*^F/F^ mice and induced *Tead1* deletion ex vivo (Fig. 4j and Supplementary Fig. S4a-b), as described above, using Ad-Cre-GFP (iACMs) or Ad-GFP (control ACMs), resulting in a significant decrease in *Tead1* transcript and protein levels (Fig. 4j and 4l). There was a small but significant decrease in the expression of MEF2d and GATA4, while ANP, BMPA, and MYH7 (β-MHC) were unchanged with TEAD1 deletion (Supplementary Fig. S4c). Consistent with that seen in iNCMs, iACMs with *Tead1* deletion demonstrated a significant reduction in mRNA expression of *NRF2* and its targets (*NQO1, HO-1, and SOD2*) compared to the GFP control ACMs (Fig. 4k) though SLC7a11, also a target of NRF2 was surprisingly increased. NRF2 and SOD2 protein levels were reduced ∼50% in TEAD1KO iACMs as compared to the control Ad-GFP treated ACMs (Fig. 4l). Assessment of the total cellular hydrogen peroxide (boronate assay) in TEAD1KO-iAMCs and control ACMs revealed that while there was a trend for an increase in H2O2 accumulation in the basal state in TEAD1KO-iACMs, exposure to 10µM DMNQ for 2h, a pharmacological inducer of ROS, elicited a significant ∼40% increase in H2O2 accumulation with TEAD1 KO, as compared to the control (Fig. 4m). Concordant with this finding TEAD1KO-iACMs accumulated significantly more (∼90%) H2O2 when challenged with 200nM AngII for 48h as compared to the Ad-GFP control ACMs, and this increase was prevented by Losartan (1uM, 48h), an AT1R blocker (Fig. 4n). These results indicate that TEAD1 function is required for normal NRF2 expression, and function, and mitigating the cellular accumulation of ROS in adult CMs under stress.

### Generation and characterization of cardiomyocyte-specific *Tead1* mosaic-deletion mice

Deletion of *Tead1* in cardiomyocytes leads to lethality, whether induced embryonically^13^ or in adult mice^14^, limiting the analysis of TEAD1 function in cardiomyocytes in vivo. To determine the cell-autonomous function of TEAD1 in cardiomyocytes in vivo, a mouse model with mosaic *Tead1* knockout in the heart (mT1KO) was generated, in which *Tead1* is deleted in only ∼50% of cardiomyocytes. *Tead1*^F/F^ mice were crossed with mTmG reporter transgenic mice to generate homozygous *Tead1*^F/F^ mice that were also homozygous for mTmG reporter transgene (mTmG-*Tead1*^F/F^). These mice were injected with AAV9 virus (2×10^9^ viral genomes/gm of bodyweight) carrying TnT-promoter driven Cre recombinase (Fig. 5a and Supplementary Fig. S5a): Group 1 was injected at postnatal day 4 (P4) which coincides with the peak cardiomyocyte proliferation window in neonatal mice (P1-P7) and assessed after 15 days for determining the function of TEAD1 in the neonatal CM proliferative period and the acute effects of its deletion. Group 2 was injected at day P4 and assessed at 5 months of age to determine the function of TEAD1 in the neonatal CM proliferative and adult CM hypertrophic growth period and the chronic effects of its deletion. AAV9-TnT-Cre injection was effective in deletion of *Tead1* in ∼50% of the cardiomyocytes when evaluated, either acutely after 15 days after injection (Group 1) or after chronic deletion for 5 months (Group 2), by western blotting (Fig. 5b & 5h) and immunostaining (Fig. 5c & 5i). This ∼50% loss of function did not lead to a change in cardiac function compared to controls as assessed by echocardiogram at the assessed time points (Fig. 5d & 5j).

**Figure 5:**
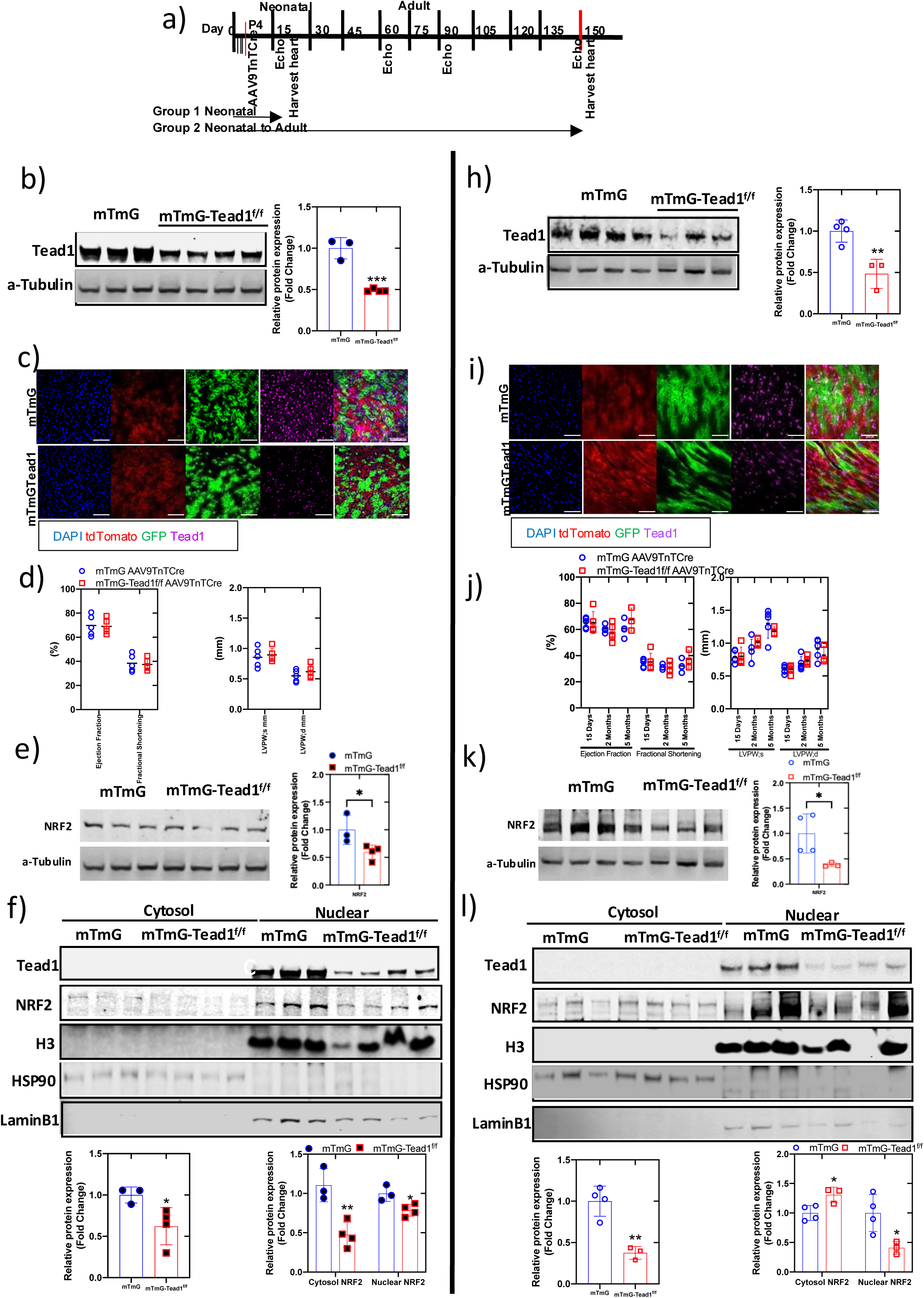
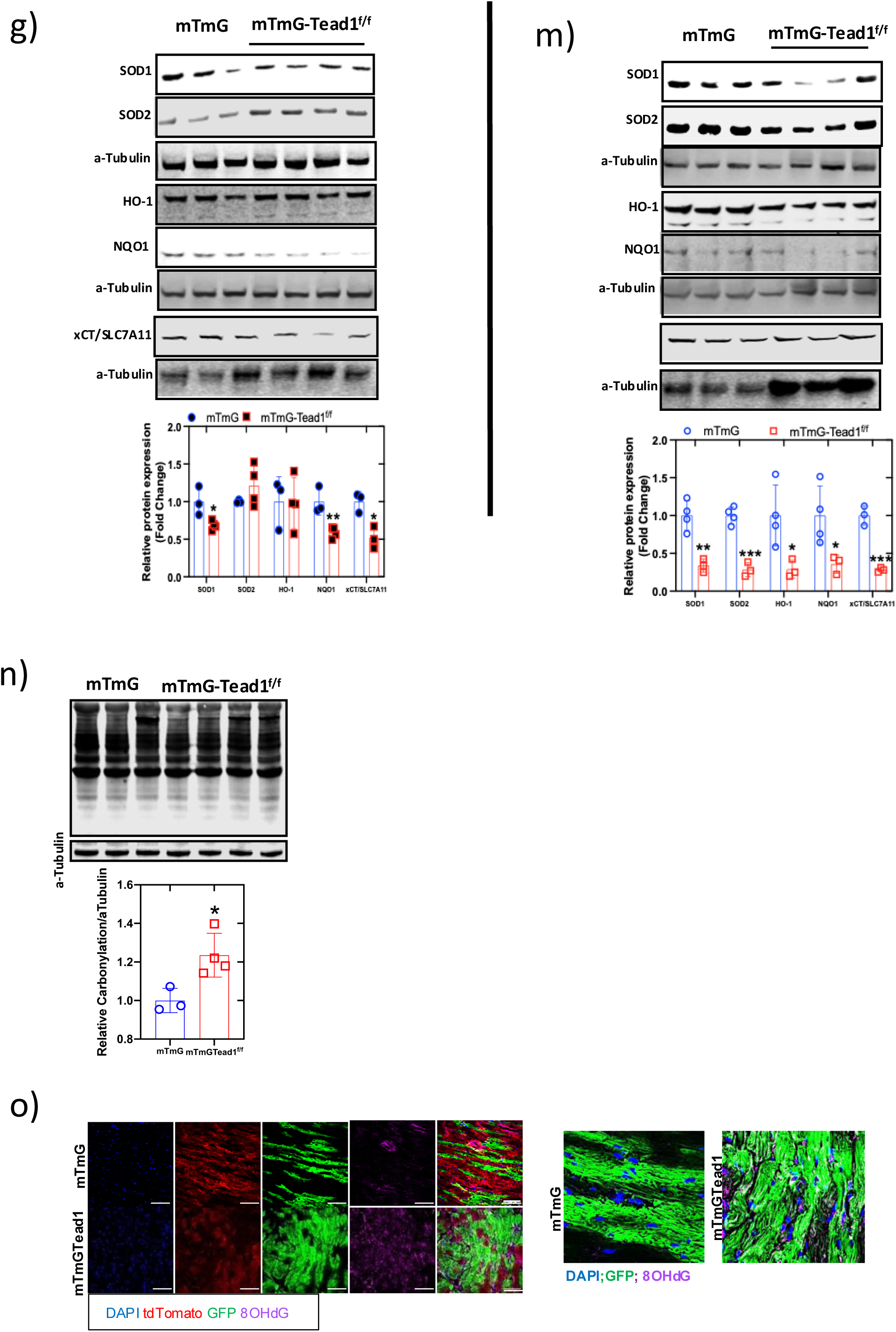
Mosaic CM-specific TEAD1 deletion leads to increased oxidative stress. **a)** Schematic of experimental timeline. Timeline to assess the effect of adult heart Tead1 mosaic knockout (T1mKO) models with AAV9TnTcre intravenous (i.v.) injection. Neonatal mTmG and mTmG-Tead1f/f (postnatal day 4-P4) mice were injected with AAV9-TnTcre (2×10^9 viral genomes/g body weight). Group 1: Cardiac function was assessed at different time points, as shown, and hearts were harvested at 15 days old [Group 1 – (b-g)] or at 5 months [Group 2 – (h-m)]. **b & h)** Immunoblotting of whole heart TEAD1 level (left) and quantification (right), with α-Tubulin as loading control. **c & i)** Representative immunofluorescent images of mosaic TEAD1 KO and control hearts. Control (mTmG) (top) and mosaic TEAD1KO (mTmG-Tead1f/f) (bottom) [GFP-green, tdTomato-red, anti-TEAD1-magenta, and nucleus-DAPI-blue staining]. **d & j)** Left ventricular (LV) function: ejection fraction (%EF), fractional shortening (%FS), LVPW;s, and LVPW;d (n=5). **e & k)** Whole heart immunoblot analysis of NRF2 in mTmG-Tead1f/f (T1mKO) and mTmG (control) (left) and quantification (right), with *α*-Tubulin as loading control. **f & l)** Nuclear fraction of whole heart immunoblot analysis of NRF2 in mTmG-Tead1f/f (T1mKO) and mTmG (control) (top) and quantification (bottom), with HsP90 for cytosol and H3 for nuclear loading controls. **g & m)** Whole heart immunoblot analysis of antioxidant proteins, SOD1, SOD2, HO-1, NQO1, and xCT/SLC7A11 in mTmG-Tead1f/f (T1mKO) and mTmG (control) (top) and quantification (bottom), with α-Tubulin as loading control. **n)** Protein carbonylation was analyzed using an anti-DNP antibody, with immunoblot of neonatal mT1KO heart whole lysate (top) and quantification (bottom). **o)** Representative immunofluorescent images of control (mTmG) and T1mKO (mTmG-Tead1f/f), showing 8-OHdG labeling in control (top) and Tead1KO (bottom) [GFP-green, tdTomato-red, anti-8-hydroxydeoxyguanosine (anti-8-OHdG)-magenta, and nucleus-DAPI-blue staining]. Statistical significance is indicated as follows: no symbol (non-significant, P > 0.05), *P < 0.05, **P < 0.01, ***P < 0.001, ****P < 0.0001, compared with control (Student’s t-test). Scale bar: 100 µm.

### Increased oxidative stress is seen with cardiomyocyte-specific TEAD1-deletion in vivo

To investigate the role of TEAD1 on oxidative stress in vivo, we analyzed the heart tissue from this mosaic cardiomyocyte-specific TEAD1 deletion mice with no systolic dysfunction. Oxidative stress occurs from a combination of an increased generation of ROS or from impaired antioxidant response. Based on our previous work^15^ and data from the in vitro studies shown above, mitochondrial dysfunction and higher ROS were seen with a loss of TEAD1 function. Here, we determined the levels of critical antioxidant proteins, and NRF2 were significantly lower with both acute and chronic loss of TEAD1 function in mosaic *TEAD1*KO hearts compared to controls (Fig. 5e & 5k). This reduction in total NRF2 resulted in a decrease in nuclear NRF2 levels (Fig. 5f & 5l). This was accompanied by a significant decrease in important antioxidant proteins such as SOD1, NQO1 and SLC7A11 in the acute mosaic *TEAD1*KO heart lysates compared to controls, while in addition to these, SOD2 and HO-1 were decreased significantly in the chronic mosaic *TEAD1*KO heart (Fig. 5g & 5m). We then assessed if changes in these antioxidant proteins resulted in oxidative stress. Protein carbonylation, a measure of oxidative stress, was significantly increased in the lysates from the left ventricles of the mosaic *TEAD1*KO and controls (Fig. 5n). 8-Hydroxy-2’deoxyguanosine (8-OHdG), an oxidative derivative of deoxyguanosine in the DNA and another indicator of cellular oxidative damage, was increased on immunostaining in the mosaic *TEAD1*KO hearts, specifically in the TEAD1-GFP-pos cardiomyocytes, as compared to controls (Fig. 5o).

### TEAD1 function is required for mitigating Angiotensin II-induced oxidative stress

Many external stimuli induce oxidative stress in cardiomyocytes, including Angiotensin II, which has been implicated in pathologic cardiac remodeling and heart failure. To test if TEAD1 function is required for mitigating Angiotensin II-induced oxidative stress in vivo, we infused Angiotensin II (1µg/kg/min) by osmotic pump for 15 days into mosaic *Tead1*KO and control mice (schematically shown in Fig. 6a). There was no change in systolic function on echocardiography between the controls and mosaic *Tead1*KO mice after Angiotensin II infusion (Fig. 6b). As expected, there was a significant increase in indexed heart weight after Angiotensin II infusion in both groups, though there was a mild but significant lower indexed heart weight in the mosaic *Tead11*KO mice, compared to control, post-Angiotensin II infusion (Fig. 6c). There was a significant increase in oxidative stress markers in mosaic *Tead1*KO mice, including 8OHdG specifically in *Tead1*-deleted green cardiomyocytes (Fig. 6d and Supplementary Fig. 6a-b) along with other markers such as lipid peroxidation and protein carbonylation in whole heart lysates (Figs. 6e-f). This was accompanied by an increase in disordered myofibers with inflammatory cells (Fig. 6g) and increased fibrosis on Masons Trichrome staining (Fig. 6h) in Angiotensin II-infused mosaic *Tead1*KO mice compared to controls.

**Figure 6:**
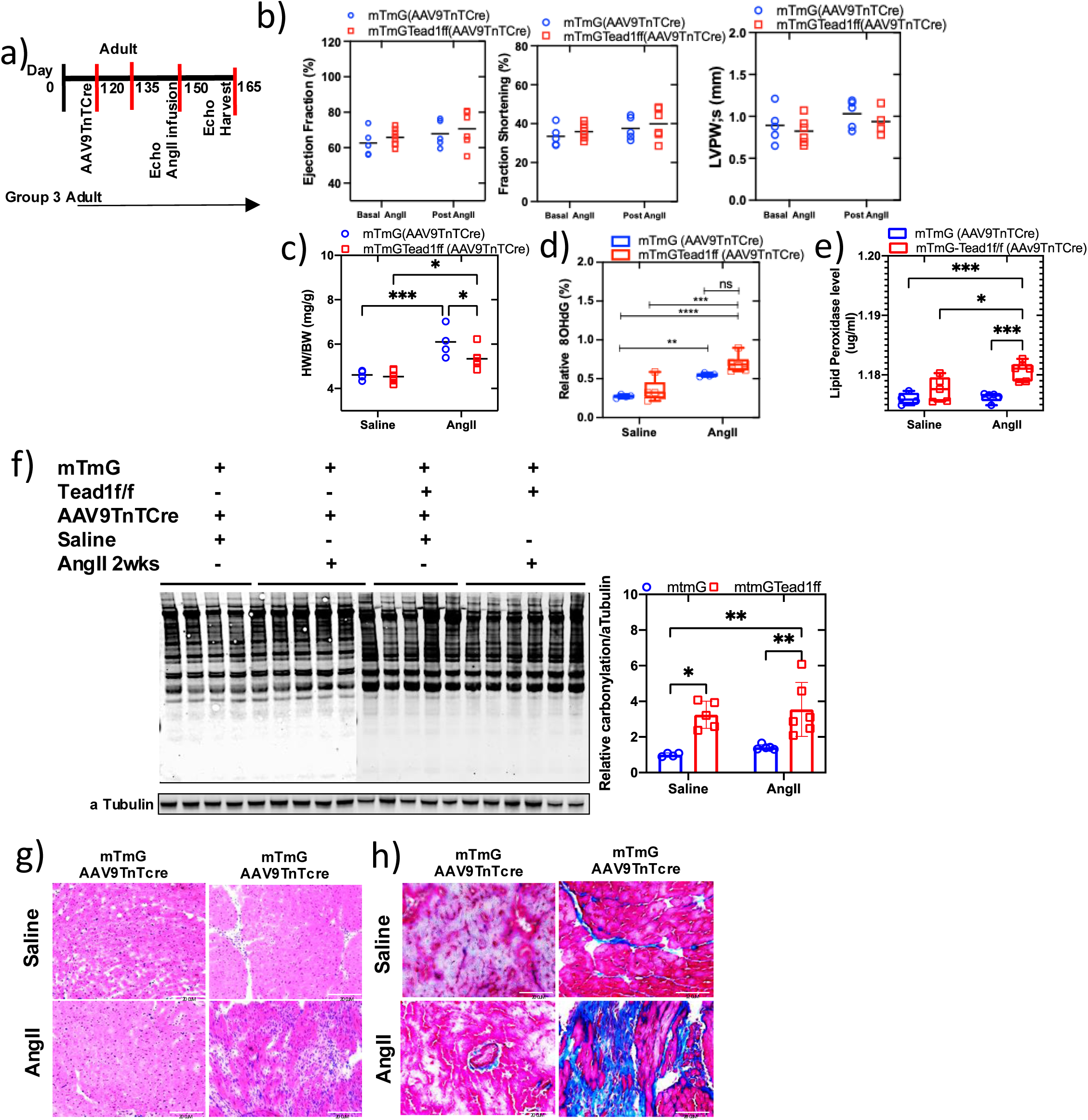
TEAD1 function is required for mitigating Angiotensin II-induced oxidative stress. **a)** Schematic of experimental timeline. Timeline to assess the effect of adult heart Tead1 mosaic knockout (T1mKO) models with AAV9TnTcre intravenous (i.v.) injection. Group 3: 3 month old adult mTmG and mTmG-Tead1f/f mice were injected with AAV9-TnT-Cre virus (2×10^9 viral genomes/g body weight). Infusion of AngII [mTmG-AAV9TnTCre (n=5) and mTmGTead1f/f-AAV9TnTcre (n=6)] or saline [mTmG-AAV9TnTcre (n=4) and mTmG-Tead1f/f-AAV9-TnTcre (n=6)] via Alzet pumps for 15 days. **b)** Ejection fraction (%EF), fractional shortening (%FS), and LVPW;d (n=4-6) and **c)** Indexed heart weight (heart weight/body weight) in mTmG-AAV9-TnTCre-AngII/saline15d (n=4-6) and mTmG-Tead1f/f-AAV9-TnTCre-AngII/saline15d (n=5-6). **d)** Oxidative-induced DNA damage by 8OHdG assay and **e)** lipid peroxidation in all four groups (n=4-6). **f)** Protein carbonylation was analyzed using an anti-DNP antibody, with immunoblot of neonatal mosaic Tead1 KO heart whole lysate (left) and quantification (right). **g-h)** Representative images of H&E (g) and Masson’s trichrome (h) stained sections of all four groups. Groups: mTmG-AAV9-TnTCre-saline15d, mTmG-AAV9-TnTCre-AngII15d, mTmG-Tead1f/f-AAV9-TnTCre-saline15d, mTmG-Tead1f/f-AAV9-TnTCre-AngII15d. Statistical significance is indicated as follows: no symbol (non-significant, P > 0.05), *P < 0.05, **P < 0.01, ***P < 0.001, ****P < 0.0001, compared with control (two-way ANOVA). Scale bar: 100 µm.

These data demonstrate that cell-autonomous loss of TEAD1 function in cardiomyocytes in vivo is required for normal levels of NRF2 and other antioxidant proteins, and a loss of TEAD1 function results in oxidative stress and cellular damage that is significantly exacerbated by external stressors such as Angiotensin II.

### Human TEAD1 (hTEAD1) restores NRF2 activity and antioxidative response in TEAD1-deficient murine cardiomyocytes

The previous experiments demonstrated that TEAD1 deficiency led to significant changes in antioxidant proteins in a cell-autonomous manner. To test if restoring TEAD1 function rescued the antioxidant response, we overexpressed h*TEAD1* using a lentiviral vector in H9C2 cells (Fig. 7a). Consistent with the data above, hTead1OE increased NRF2-ARE reporter luciferase activity as compared to vector control (Fig. 7b). Exposure for 24 hrs to 10µM DL-sulforaphane, a Keap1 inhibitor that stabilizes NRF2 protein, led to an increase in ARE-luc activity in control cells as expected, however, with hTEAD1 OE, there was a further increase in ARE-luc activity, indicating that hTEAD1 OE was sufficient to increase NRF2 activity (Fig. 7c) significantly. hTEAD1 OE was also sufficient to increase ARE-luc activity in TEAD1-KO cells, and this was further increased by sulforaphane, indicating this is due to a TEAD1 OE-mediated increase in NRF2 protein levels (Fig. 7d). Consistent with this, while TEAD1-KO H9C2 cells displayed increased Mitosox, overexpression of hTEAD1 was sufficient to restore Mitosox to that of control cells (Fig. 7e). Immunoblotting and immunofluorescent staining analysis in neonatal cardiomyocytes isolated from Tead1-Flox mice revealed that lentivirus-mediated hTEAD1OE was sufficient to restore NRF2 protein (Fig. 7f-g). These restoration experiments support the direct role of TEAD1 in positively regulating NRF2 levels and activity in cardiomyocytes.

**Figure 7:**
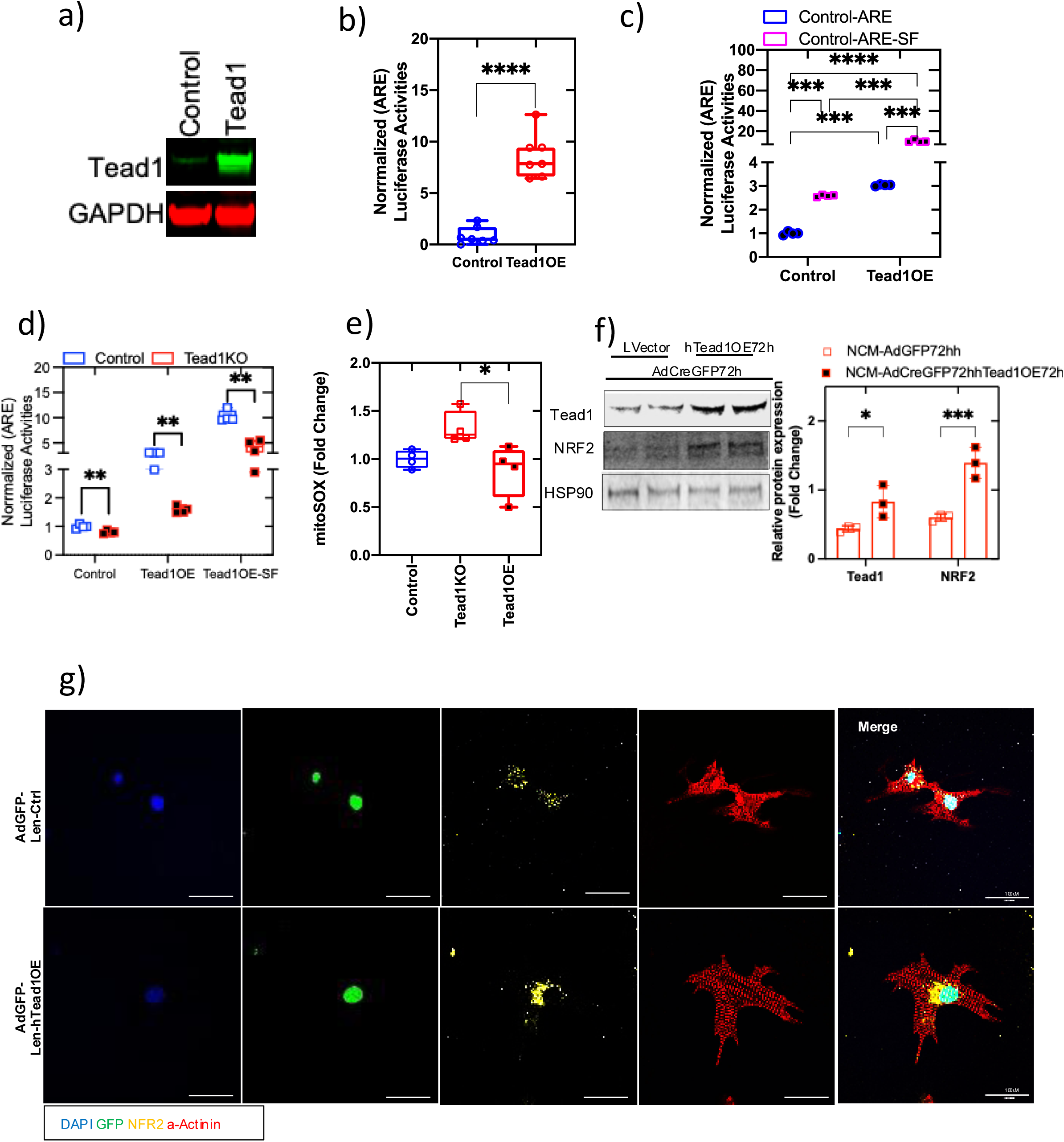
hTEAD1 restores NRF2 and antioxidant response in TEAD1-deficient murine CMs. **a)** Immunoblotting for TEAD1 protein in H9C2 cells after transduction with lenti-GFP (control) or lenti-GFP-hTEAD1. **b)** ARE-luciferase reporter assay to assess NRF2 activity in H9C2 cells 72 hours after overexpression of hTEAD1 (lenti-GFP-hTEAD1 OE) compared to control (lenti-GFP). **c)** ARE-luciferase reporter assay to assess NRF2 activity in H9C2 cells with and without 10 µM DL-sulforaphane for 24 hours. **d)** ARE-luciferase reporter assay with and without 10 µM DL-sulforaphane for 24 hours after overexpression of hTEAD1 in Tead1KO and control H9C2 cells. **e)** Quantification of superoxide levels using the mitoSOX assay in H9C2 cells with TEAD1KO, overexpression of hTEAD1 (lenti-GFP-hTead1), TEAD1OE) compared to control (lenti-GFP). **f)** Immunoblotting for TEAD1 and NRF2 in ex vivo TEAD1-depleted neonatal CM (NCM) (TEAD1 flox mouse NCM isolated and infected ex vivo with Ad-Cre virus). After 72 hours of Tead1KO, hTEAD1 (or GFP control) was overexpressed. Quantification is shown on the right, with HSP90 protein levels used as loading control. **g)** Representative immunofluorescent images of anti-NRF (yellow), GFP (green), and nucleus-DAPI (blue) staining in NCM with control virus (top) and lenti-GFP-hTEAD1 OE (bottom). Statistical significance is indicated as follows: ***p<0.0005; **p<0.005; *p<0.05; no symbol indicates no significance, compared with control (Student’s t-test and Two-way ANOVA). (n = 3–6 per group). Scale bar: 100 µm.

### TEAD1 is required for NRF2 function and antioxidant response in human iPSC-derived cardiomyocyte (hiPSC-CM)

Results from in vivo experiments in mice and in vitro experiments in rat CM cell lines demonstrated that loss of TEAD1 function resulted in transcriptional downregulation of the NRF2 pathway leading to significant oxidative stress and adverse cardiac remodeling in hearts under stress. This was consistent with human heart RNA-seq data demonstrating the downregulation of TEAD1 in heart failure and enrichment of NRF2-oxidative stress pathways (Fig.1 and Fig. S1). To confirm if TEAD1 function was similarly required in human CMs and to validate the TEAD1-regulated antioxidant response, we first established two hiPSC lines using sgRNAs directed against exon 3 of TEAD1 using a CRISPR-Cas9 system and a loss of TEAD1 was confirmed by western blotting, (Fig. 8a). After 12 days of directed differentiation into iPSC-Cardiomyocyte (hiPSC-CM) (see Methods), gene expression analysis revealed that while control iPSC-CMs displayed robust expression of CM-specific markers *MYH6, MYH7 and TNNT2*, these were expressed in TEAD1 KO iPSC-CMs, but at a significantly lower level as was *CYR61*, a known TEAD1 target gene (Fig. 8b).

**Figure 8:**
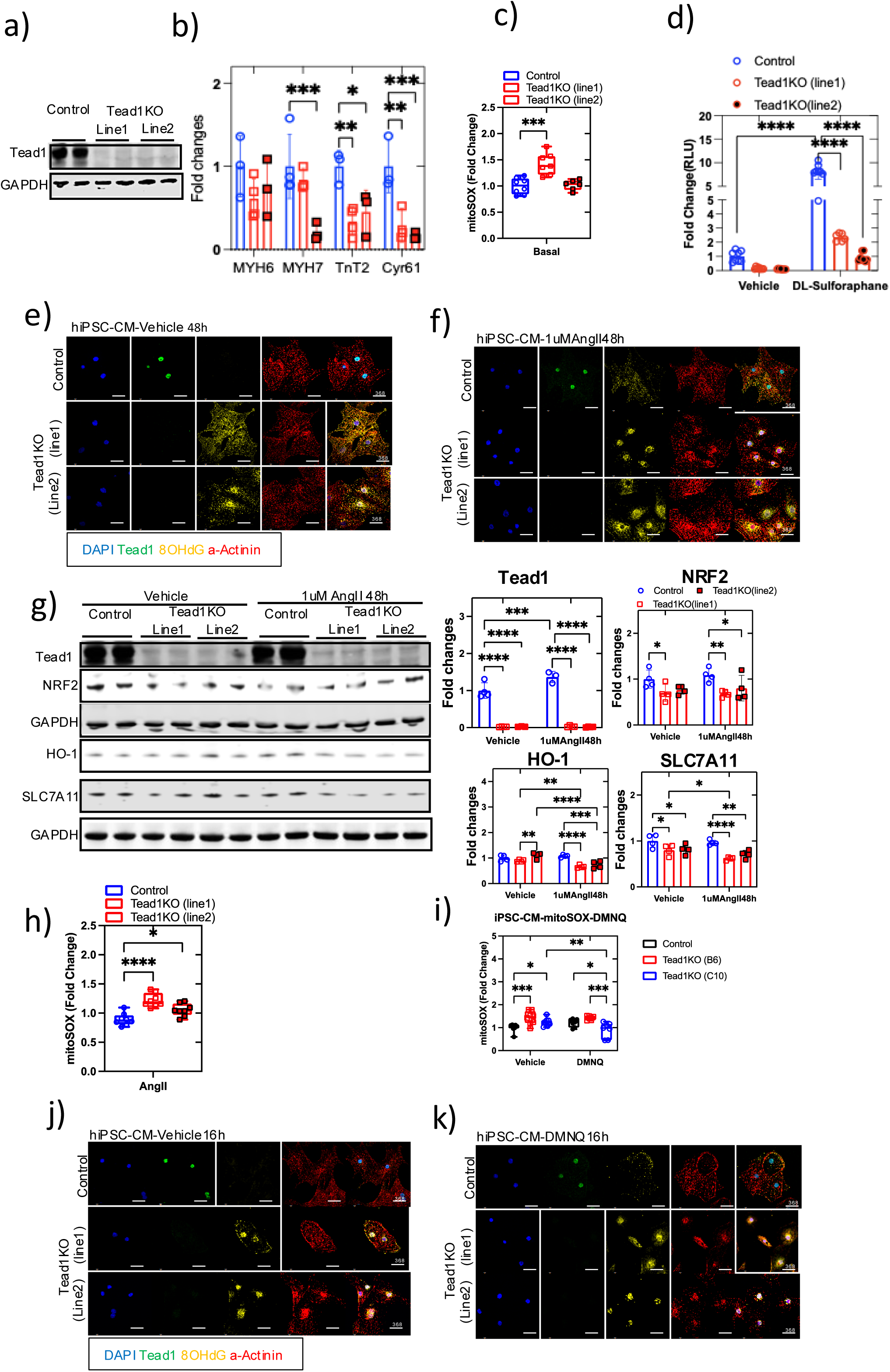
TEAD1 is required for NRF2 function and antioxidant response in hiPSC-CM. **a)** Validation of Tead1 knockout (KO) in stable iPSC-CM lines (Tead1KO-line 1 and line 2) and control cell lines, by Western immunoblotting for TEAD1 protein in whole iPSC-CM lysates, with GAPDH as a loading control. **b)** Expression of cardiomyocyte markers and TEAD1 targets (*MYH6, MYH7, TNNT2, CYR61*) in hiPSC-CM TEAD1KO and control lines assessed by qPCR normalized to housekeeping genes expression. Fold change over control lines is shown. **c)** Quantification of superoxide levels using the mitoSOX assay in hiPSC-CM TEAD1KO and control lines. **d)** ARE-luciferase reporter assay to assess NRF2 activity in basal and 10 µM sulforaphane-treated (16 hours) hiPSC-CM TEAD1KO and control lines. **e-f)** Representative immunofluorescent images of vehicle- (**e**) or AngII- (**f**) treated (48 hours) hiPSC-CM TEAD1KO and control lines, showing 8-OHdG labeling in control (top) and TEAD1KO (bottom) [anti-TEAD1 (green), anti-8-hydroxydeoxyguanosine (anti-8-OHdG) (yellow), anti-alpha actinin (red), and nucleus-DAPI (blue) staining]. **g)** Immunoblotting for TEAD1, NRF2, HO-1, and SCL7A11 protein levels in whole hiPSC-CM lysates of TEAD1KO and control lines, with GAPDH as a loading control and quantification shown on the right. **h)** Quantification of superoxide levels using the mitoSOX assay in 1 µM AngII-treated (48 hours) hiPSC-CM TEAD1KO and control lines. **i)** Quantification of superoxide levels using the mitoSOX assay in DMNQ-treated (16 hours) hiPSC-CM TEAD1KO and control lines. **j & k)** Representative immunofluorescent images of DMNQ and vehicle (16 hours) treated hiPSC-CM TEAD1KO and control lines, showing 8-OHdG labeling in control (top) and TEAD1KO (bottom) [anti-Tead1 (green), anti-8-hydroxydeoxyguanosine (anti-8-OHdG) (yellow), anti-alpha actinin (red), and nucleus-DAPI (blue) staining]. Statistical significance is indicated as follows: ***p<0.0005; **p<0.005; *p<0.05; no symbol indicates no significance, compared with control (Student’s t-test and Two-way ANOVA). Scale bar: 36.8 µm. (n = 3–8 per group).

Examination of the effect of the loss of TEAD1 function on oxidative stress revealed that under basal conditions, the superoxide levels (Mitosox assay) were elevated in the TEAD1KO cell line (Fig. 8c), similar to what was seen in the in vitro and in vivo mouse studies. Furthermore, a significant decrease in ARE-luc activity was seen in TEAD1 KO hiPSC-CMs, as compared to controls and the increase with sulforaphane exposure seen in control cells was significantly diminished with TEAD1 KO in hiPSC-CMs (Fig. 8d). This was accompanied by an increase in oxidative stress-induced DNA damage, indicated increased by 8OHdG immunostaining in the Tead1KO hiPSC-CM cell lines compared to the control line, under basal conditions and was further increased after exposure to 1µMAngII for 48h (Fig. 8e-f).

The decrease in protein level of NRF2 and its targets were decreased in TEAD1 KO lines both in basal and after exposure to 1uMAngII for 48h (Fig. 8g). This resulted in an increased accumulation of mitochondrial superoxide in TEAD1KO hiPSC-CMs compared to controls, after exposure to 1µMAngII for 48h (Fig. 8h). Consistent with this, exposure to DMNQ, an inducer of oxidative stress via accumulation of superoxide and hydrogen peroxide, led to increased mitosox staining in TEAD1 KO hiPSC-CMs as compared to controls (Fig. 8i) along with increased 8OHdG immunostaining under basal (Fig. 8j) and DMNQ stimulation (Fig. 8k). These results demonstrated that TEAD1 was required for normal antioxidant response in basal and stressed states in human iPSC-CMs.

## Discussion

Heart failure has been defined as “a complex clinical syndrome that can result from any structural or functional cardiac disorder that impairs the ability of the ventricle to fill with or eject blood” and affects 1 in 5 Americans^25^. Heart failure most commonly results from a failure of homeostatic pathways in myocardial cells, including cardiomyocytes, to prevent or mitigate cellular stress. This culminates in cardiac dysfunction and subsequent inability to meet the body’s perfusion demands. Oxidative stress is a frequent accompaniment and contributor to the pathophysiology of heart failure and results from an imbalance in the generation and elimination of ROS, which causes tissue damage and myocardial dysfunction^26,27^. Mitochondrial dysfunction, a frequent accompaniment of myocardial dysfunction, leads to further increases in ROS generation and aggravating oxidative stress, resulting in a vicious cycle that, along with other factors, culminates in heart failure. There is reported evidence of a decreased abundance of antioxidant enzymes and increased oxidative injury in human heart failure^28^. Recent literature has shown that oxidative stress is present in patients with early-stage cardiac failure and preserved heart function^29^. Here, we show that in human hearts, both in control and in heart failure, the expression of NRF2, the master regulator of antioxidant response, and other antioxidant genes are significantly positively correlated to the expression of TEAD1. This is consistent with the TEAD1 occupancy in the promoter regions of these genes, which also were open chromatin regions, in TEAD1-Chip-seq and ATAC-seq studies in adult mouse hearts, strongly suggesting direct transcriptional regulation of these genes by TEAD1.

The current study demonstrates the critical regulatory role of TEAD1 in mitigating ROS in cardiomyocytes in gain- and loss-of-function studies in vitro and in vivo loss-of-function studies. In addition, evidence of oxidative stress in the form of DNA damage with positive 8-hydroxy guanidine and protein carbonylation was significantly increased with TEAD1-depletion. These findings are supported by other complementary studies that have reported protection against ROS with hippo pathway modulation^30^, as in cardiac-specific MST1 knockout mice^31^, while MST1 transgenic cardiac overexpression led to DCM with severe mitochondrial dysfunction and increased ROS production with increased protein levels of NADPH oxidases, NOX2 and NOX4^32^. Similarly, in other tissues, TAZ has been shown to mitigate ROS production, such as in microglia28 and ovarian granulosa cells29; YAP has also been shown to mitigate oxidative stress in cochlear hair cells^33^ and small cell lung cancer model^34^, though the dependency on TEAD1 was not demonstrated. This is relevant as YAP binds to other transcription factors, such as Pitx2, to regulate oxidative stress, as was shown in primary and iPS-derived cardiomyocytes^35,36^.

The hippo-TEAD pathway is critical in cardiac development and adult heart function. While embryonic TEAD1 deletion in mice leads to poor cardiomyocyte proliferation, apoptosis, and embryonic lethality^13^, adult cardiomyocyte-specific deletion of TEAD1 leads to acute DCM^14^. Similarly, YAP/TAZ deletion have significant heart failure and mortality^9^, while knockout of the upstream inhibitory hippo components (SAV, MST, LATS2) or overexpression of YAP all led to increased cardiac hyperplasia^10^. However, TEAD1 deletion in adult cardiomyocytes leads to severe acute heart failure with accompanying mitochondrial dysfunction^15,16^. Hippo pathway has also been shown to play a critical role in cardiac adaptation to pressure overload, with YAP shown to mediate cardiac hypertrophy in response to pressure overload^37–39^, while LATS loss-of-function either genetically^40^ or by small molecules^41^ mitigated adverse cardiac remodeling after pressure overload. Cardiomyocyte-YAP also improved cardiac function and survival post-MI^12^. While these reports are strong evidence for the critical role of hippo pathway components in cardiac adaptation, the role of TEAD1 itself has not been studied before, and it is unknown as to what component of the compensatory response is TEAD1-dependant, especially the cellular stress responses, such as oxidative stress, which are critical components of cardiac remodeling. Mitigating oxidative stress would be critical for any protective response during cardiac remodeling. Here, we demonstrate that TEAD1 is a critical regulator of the antioxidant stress response in cardiomyocytes, both in the basal state and during Angiotensin II-induced cardiac remodeling, by directly regulating the levels of NRF2 and other antioxidant enzymes in a cell-autonomous manner. This is consistent with the analysis presented here of the significant correlation between the decreased expression of TEAD1 and NRF2 during human heart failure seen in RNA-seq datasets, making the TEAD1-NRF2-antioxidant response axis a potential therapeutic target for mitigating adverse cardiac remodeling.

TEAD1 is a critical cardiac transcription factor that interacts with many others, including GATA4, NKX2-5, MEF2A, MEF2C, SRF, and TBX5, in a combinatorial manner to regulate cardiomyocyte specification and adaptation^42–44^ by controlling core cardiomyocyte primary functions. Data from our adult wild-type (WT) mice endogenous (chromatin immunoprecipitation sequencing) TEAD1-ChIP-seq analysis revealed that TEAD1 bound to the promoter regions of many critical pathways, in addition to antioxidant NRF2 pathway, consistent with those reported earlier^43–45^. While many of the pathways enriched for TEAD1 occupancy in their promoter regions are very relevant to cardiac adaptation, a detailed assessment of all these pathways in TEAD1 mosaic knockout is beyond the scope of this study and will be the subject of future studies. Interestingly, the AngII challenge for 15 days led to a significant increase in fibrosis and inflammatory cells, suggesting activation of profibrotic and proinflammatory signaling pathways in the mosaic TEAD1 KO mice hearts. This adverse cardiac tissue remodeling, when only 50% of cardiomyocytes have TEAD1 loss-of-function, indicates the critical nature of TEAD1 in maintaining normal cardiac cellular milieu under conditions of neurohumoral stress. In addition to an increase in oxidative stress with Tead1-depletion, there are likely to be other pathways affected in the TEAD1 regulated signalome in cardiomyocytes and cross talk with non-cardiomyocyte myocardial cellular population, and these will also be the subject of future studies.

Our findings demonstrated that loss of TEAD1 function in human iPSC-CMs led to a significant downregulation of NRF2 activity and a consequent increase in superoxide levels. In hiPSCs, ROS generation was substantially increased with the onset of reprogramming, and decreasing ROS levels via antioxidants or NOX inhibitors significantly decreased reprogramming efficiency^46^. In support of our current study, iPSC-CM from patients with HLHS showed that early heart failure (HF) is associated with increased apoptosis, mitochondrial respiration defects, redox stress from abnormal mitochondrial permeability transition pore (mPTP) opening, and failed antioxidant response, with no translocation YAP and NRF2 into the nucleus. However, iPSC-CM from patients without early HF showed normal respiration with an elevated antioxidant response, with translocation of YAP and NRF2 into the nucleus^47^. This also raises the possibility of whether YAP-TEAD1 had a role in this NRF2-mediated redox response. Nuclear translocation of YAP was significantly lower in Duchenne muscular dystrophy (DMD-iPSC-CMs); the dystrophic heart may contribute to DMD-cardiomyopathy pathogenesis^48^. Increased nuclear 8OHdG immunostaining with 1uMAngII 48h treatments in the iPSC-CM Tead1KO cell lines supports our other in vivo and in vitro analyses. These support the critical regulatory role of TEAD1 and the mammalian hippo pathway in preserving the antioxidant response and mitigating oxidative stress-induced cardiac dysfunction.

In conclusion, this study demonstrates that TEAD1 is a direct transcriptional activator of NRF2, regulates ROS in cardiomyocytes, and is critical in mitigating oxidative damage during Angiotensin II-induced hypertrophy and pathological cardiac remodeling. This TEAD1-NRF2 regulation is conserved in both mouse and human cardiomyocytes. Targeting this axis and TEAD1 could be a promising therapeutic strategy for preventing and treating cardiac dysfunction.

## Methods

### Mosaic Tead1knockout heart and echocardiography analysis

Animal studies were performed per the ethical standards of the University of Pittsburgh (Pitt) under an approved Institutional Animal Care and Use Committee (IACUC) protocol. In the present study, male C58Bl6J mice were used. Animals were housed in a standard temperature-controlled environment under a 12-h light-dark cycle. Tead1 flox/flox mice (Tead1f/f homozygous) were crossed with mTmG mice (B6.129(Cg) Gt(ROSA)26Sortm4(ACTB-tdTomato,-EGFP)Luo**/**J), created Tead1f/f-mTmG homo mice were used for experiments. Tead1f/f-mTmG (for Tead1KO) and mTmG (for Control) P4 pups and adult mice P120 were tail vein injected with 2×109/g and 5×109/g AAV9TnTCre respectively, and post 15 days (Group1), 5months (Group 2&3), were subjected for cardiac measurements by echocardiography. Group 1 and 2 mice were sacrificed, and the heart was harvested for histological, protein, and gene expression analysis. Group 3 mice were treated with 1µg/kg/min Angiotensin II (AngII) /Saline alzet pump (Model # 1004) infusion for 15 days, and mice were further assessed for cardiac function post 15 days of AngII infusion. Mice were sacrificed and heart processed for histological, protein, and gene expression analysis.

Echocardiography was performed on mice with the Vevo2100 machine (VisualSonics, Toronto) and an MS550D transducer. Mice were anesthetized by isoflurane (3% for induction, 1.7% for experiment), and body temperature was monitored using a rectal temperature probe and controlled at 36-37 °C.

### Ex vivo Cardiomyocyte Tead1 Knockout Transcriptome Analysis

Neonatal Cardiomyocytes (NCMs) were isolated, according to the manufacturer protocol (Pierce Primary Cardiomyocyte Isolation Kit #88281Y), from Tead1^Flox/Flox^ P1-2 old pup hearts. They were then infected with Adenovirus GFP (control) or GPF-Cre (Tead1KO) at 100 MOI, and total RNA extracted after 72hr, and RNA-seq was performed was done by Novogene. cDNA was generated using Clontech SMART-seq v4 and libraries using Nextera XT (Illumina FC-131– 1024), and samples were sequenced at 125 bp, ∼40 million paired-end reads/sample, on Illumina HiSeq 2500. All samples were subjected to a quality control assessment using the FASTQC package. All the files of RNA-seq reads were aligned to the GRCm38 (mm10) using STAR with default settings. Gene counts were obtained using the feature count utility of the subread five packages. The count data were then rld (rlog transformed counts)-normalized (the rld function transforms the count data to the log2 scale to minimize differences between samples for rows with small counts and normalized to library size). Differential expression analysis was performed using the DESeq2 R package; we identified markers as those genes with an FDR-corrected p-value < 0.05 and a log2 (fold change) > 0.58. Functional enrichment analyses were performed using GSEA and IPA. All RNA-seq tests were FDR-adjusted for multiple testing corrections.

### Publicly available RNA seq data analysis

Publicly available data^17–20^ were processed separately from the methods mentioned earlier. In addition, the analysis was validated with GREIN^48,49^.

### Chromatin Immunoprecipitation sequencing (ChIP-seq) and analysis

Wild-type C57Bl6 mice hearts (12 weeks old) were removed, flash frozen, and sent for Tead1-ChIP seq done by Active Motif (Carlsbad, CA). Raw sequence reads were aligned to the GRCm38 (mm10) reference assembly using Bowtie2 v2.4.2. Peaks were defined using MACS2 with FDR (p-value cutoff 0.05). Using GREAT (4.0.4), each gene was annotated - a basal regulatory domain of a minimal distance of 5 kb upstream and 1 kb downstream of the transcription start site (TSS), and an extended regulatory domain up to 1000 kb upstream and downstream of the gene’s basal domain. The MEME tool suite (5.4.1) was used for de novo TF motif discovery.

### Ingenuity Pathways Core Analysis Search Parameters

For IPA core analysis of the RNA seq (Fold Change of 1.2 and p value<=0.05), ChIP-seq (log2 fold enrichments ≥5) with an ATAC seq, uploaded data fields included the gene name and log2 value for gene names and used the default setting.

### Adult cardiomyocyte Isolation

Adult cardiomyocytes (ACM) were isolated from 8-12 old mice weeks, as previously described^14^, and were used in downstream analysis such as ROS assays (Coumarin Boronate and MitoSox assay), gene expression (Realtime qPCR), immunoblotting, and immunostaining.

### Gene Expression validated by real-time qPCR

Total RNA was isolated from cell lines via Trizol extraction using a standard protocol with DNase treatment (ZYMO RNA isolation Kit #R-2072), and cDNA was generated using the cDNA synthesis Master Mix (GenDEPOT # R6200-500). RT-qPCR reactions were run in qPCR master mix (GenDEPOT #Q5603-005 Real-time qPCR machines (QuantStudio-Applied Biosystem), and Ct values were normalized to the 18sRNA, GAPDH, B2M and PPIA housekeeping gene.

### CRISPR-Cas9 knockout and overexpression studies

CRISPR-Cas9 gene-editing experiments were carried out using the lentiCRISPRv2 lentiviral system (Addgene, 52961). Guide RNAs (gRNAs) targeting exon 3 of TEAD1 were designed using CRISPR Design and cloned into the lentiCRISPRv2 vector. The lentiCRISPR v2 vector with a gRNA insert (5000 ng), the packaging plasmid psPAX2 (3750 ng, Addgene #12260), and the envelope plasmid pMD2.G (1250 ng, Addgene #12259) were mixed and then added to a mixture of 60 μl X-tremeGENE 9 DNA Transfection Reagent (Roche) and 1 ml OPTI-MEM (Thermo Fisher Scientific, 31985070). 293-FT cells were used for transfection and harvesting of the lentivirus at 72 h. Then, cells were infected at 50% confluency with freshly collected CRISPR-Cas9-gRNA lentivirus supplemented with 8 μg ml−1 polybrene (Sigma-Aldrich, TR-1003-G). Infected cells were selected in media using puromycin treatment for up to 14 days (2 μg ml−1). Scramble gRNA used as a control cell ensured uniform cell death within 14 days of puromycin treatment. For overexpression studies, the (GFP-tagged human TEAD1 (Lenti-GFP-hTEAD1) and (Lenti-GFP backbone) lentivirus (Applied Biological Materials Inc., #BC115398). Infected cells were selected in media using puromycin treatment for up to 14 days (2 μg/ml). Puromycin concentration was determined by 100% uniform cell death within 14 days of puromycin treatment in control cells.

### Western immunoblot and immunofluorescence

We have used the protocol for tissue lysis and subcellular fractioned^50^ with some modification in homogenize and fractionation buffer [0.25M sucrose; 1mM EDTA,3mMHEPES and Halt (ThermoFisher #1861281) pH 7.2]. All the in vitro and ex vivo cells were lysis with NP40 buffer (Invitrogen # FNN0012) and Halt (ThermoFisher #1861281). Immunoblotting was performed by standard protocol using the following antibodies: TEAD1 (Cell signaling #12292S, 1:1000, abcam #133533, 1:1000), NRF2 (Cell signaling #12721S, 1:1000),SOD1 (Santa Cruz Biotech# sc-101523, 1:1000), SOD2 (Abcam, ab137037, 1:1000), HO-1 (Cell signaling #86806S, 1:1000), NQO1 (Cell signaling #3187S, 1:1000), xCT/SLC7A11 (Cell signaling #12691S, 1:1000), 8OHdG (DR1001, 1:200-for IF), Cre (Cell signaling #15036S, 1:500-for IF), HSP90 (Santa Cruz Biotech# sc-13119, 1:1000), a-Tubulin (Proteintech# 66031, 1:10000), Lamin B1 (Proteintech# 66095, 1:100000), a-Actinin (Sigma# A7732, 1:500-IF) and H3 (Cell signaling #4499S, 1:2000), which were diluted in 2.5% BSA (or) 5% nonfat-milk in TBS-T solution and incubated for overnight at 4 °C, followed by species-appropriate fluorochrome-conjugated secondary antibody anti-Rabbit DyLight 800 (Cell signaling #5151S, 1:5000), anti-mouse DyLight 680 (Cell signaling #5470S, 1:5000), to detect immunoreactivity with LI-COR instrument.

Immunofluorescence analysis was performed on 8 micron cryosectioned mouse heart tissue. After resection, the mouse heart was perfused with 2% paraformaldehyde (PFA) for 1 h, followed by 30% sucrose overnight before being frozen in Isopentane for 30 seconds and then stored at -80 degrees C. All sections underwent: blocking in 10% normal donkey serum (NDS)/0.5% triton-X (TX) for 1 h at RT; primary antibody incubation (1% NDS/0.25% TX overnight at 4 °C) with TEAD1 (Cell signaling #12292S, 1:200), 8OHdG (DR1001, 1:200-for IF), Cre (Cell signaling #15036S, 1:500-for IF), a-Actinin (Sigma# A7732, 1:500-IF); and species-appropriate fluorochrome-conjugated secondary antibody, incubation in 1% NDS/0.25% TX for 1-2 h at RT. Nuclear counterstain was done with DAPI (UPitt imaging core provided). Images were captured on a confocal Nikon and Leica Stellaris 5 confocal microscope.

### In-vitro and ex-vivo cell imaging

Cells were plated in 96well (Boronate assay, mitoSox assay) 6-12 well plate (protein/RNA) and 6 well plate coverslip/chamber slide coated with gelatin/fibronectin (H9C2 cell study) laminin (or) Geltrex (primary cardiomyocyte) overnight 4 °C and washed in PBS. Cells were plated and subjected to the following treatment conditions [1µMAngiotensin II 48h/vehicle (PBS); 10mM DNMQ 16h or Vehicle (DMSO); 10µMAntimycin A 1h/Vehicle (DMSO); 200µM Hydrogen peroxide 2h/Vehicle (media); 10uMLosartan 48h/Vehicle (water) and 10uM DLSulforaphane 24h/Vehicle (DMSO)], according to the experiments and after washing three times with 1xPBS, samples were fixed with 2% paraformaldehyde (PFA) and washed with 1xPBS. Cells were permeabilized with 0.1% Triton X-100 made in PBS solution for 15 min and washed 4x with PBS+0.5% BSA (PBB). The samples were blocked with PBS+ 2% BSA for 1h and washed with PBB. Samples were incubated in primary antibody TEAD1 (Cell signaling #12292S, 1:200), 8OHdG (DR1001, 1:200-for IF), NRF2 (Cell signaling #12721S, 1:100), diluted in PBB for 1h and Washed 3X with PBB. Samples were incubated in species-appropriate fluorochrome-conjugated secondary antibody in PBB for 1h and washed three times with PBB, followed by DAPI stain for 30 seconds and after PBS washes, mounted with mounting media with coverslips and imaged in Nikon confocal.

### Reactive oxygen species and mitochondrial superoxide quantifications

Total cellular H2O2 as an indicator of reactive oxygen species (ROS) generation was quantitated by fluorescence-based Coumarin boronate acid (CBA) assay, as per manufacturer’s protocol (Cayman Chemical). Mitochondrial superoxide estimated by mitoSOX assay according to manufacture (Invitrogen # M36008) final step used measurement at without phenol red condition. Cells were plated in 96well (Boronate assay, mitoSox assay) coated with gelatin/fibronectin (H9C2 cell study) Laminin (primary cardiomyocyte) overnight 4 °C and washed in PBS. Cells were plated and follow the treatment conditions [(1uMAngiotensin II48h/vehicle (PBS); (10mMDNMQ16h/Vehicle (DMSO); (10uMAntimycin A1h/Vehicle (DMSO); (200uM Hydrogen peroxide 2h/Vehicle (media) ; (10uMLosartan48h/Vehicle (water) and (10uM DLSulforaphane24h/Vehicle (DMSO)], according to the experiments.

### Biochemical measurement of lipid peroxides, oxidative DNA damage, and protein carbonyl content

Oxidative DNA damage accumulation in the heart was detected by measuring 8-hydroxydeoxyguanosine by ELISA, as per manufacturer’s protocols (Cell Biolabs, Inc. #STA-320). Lipid peroxide accumulation in the heart was detected by measuring Lipid Peroxidation (4-HNE) (abcam #ab238538). The lipid peroxide level in the heart is expressed in micrograms per milliliter, and the oxidative DNA Damage level in the heart is expressed as a percentage of total nuclei.

Protein carbonyl content was assayed using the Oxyblot kit (Millipore, #S7150). Whole heart tissue and cell lysate were obtained as described above. Protein content was assayed with BCA assay. 2,4-Dinitrophenylhydrazine was added to samples to derivatize carbonyl groups from the protein side chains. Derivatized samples were then separated using electrophoresis, as described above. Western blot analysis was carried out using the 2,4-dinitrophenylhydrazine antibody provided (1:150). Densitometry was carried out using the integrated intensity value for each band. The total densitometric value of all bands within each lane was used to measure protein carbonyl content overall. Analyses of results were carried out as a ratio of protein to alpha-tubulin.

### Tead1 and Nrf2-ARE activity assays

Luciferase assays were performed in H9C2 cells (scramble control and TEAD1KO cells) with specified treatment conditions. To measure TEAD1 activity, cells in 12 or 24 plates were co-transfected overnight with 8xTEAD-Luc (0.25 µg cm−2) plus the indicated DNA constructs: Lenti-GFP, Lenti-TEAD1-OE. To measure NRF2-ARE activity, H9C2/Neonatal Cardiomyocyte cells (scramble control, TEAD1KO cells, TEAD1-OE) with specified treatment conditions. NRF2-ARE measurement was performed according to the manufacturer protocol (BPS Bioscience,#60514). Luciferase normalization was performed in every case by co-transfecting a Renilla Luciferase Vector (Promega). After 48 hours, luciferase activity was measured using a Dual-Glo Luciferase Assay Kit (Promega) and a Microtiter plate luminometer (GloMax Navigator, Promega).

### Statistics

All analyses were performed in triplicate or greater, and the means obtained were used for ANOVA or independent t-tests. Statistical analyses, variation estimation, and validation of test assumptions were carried out using the Prism 7 statistical analysis program (GraphPad). (\***p** < 0.05, \*\***p** < 0.01, \*\*\***p** < 0.001), unless otherwise indicated.

## Acknowledgments

This work was supported by grants from the National Institutes of Health R01-HL147946 (M.M.), R01 DK130499 and R01DK128972 (V.Y.), and the VA-ORD - I01BX002678 (V.Y.).

