## Supplementary Figures for "TEAD1 is a novel regulator of NRF2 and oxidative stress response in cardiomyocytes"

#### Supplementary Figure 1

a

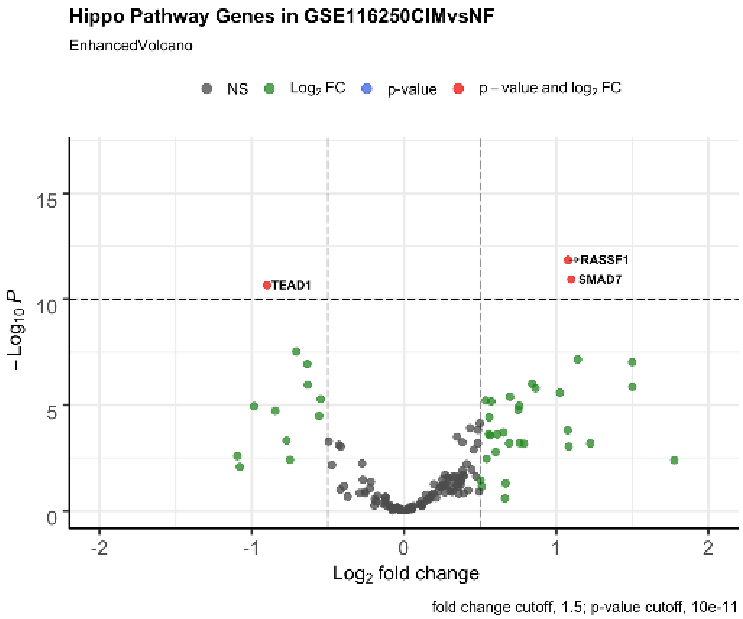


b

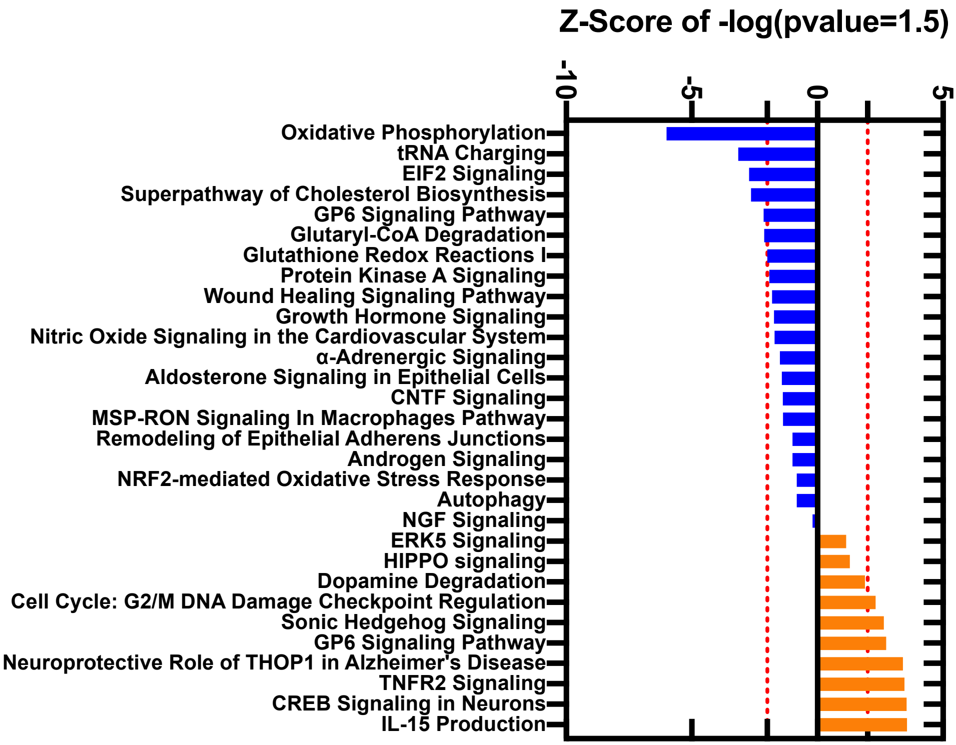

**Supplementary Figure 1: TEAD1 expression is positively correlated with antioxidant gene expression in human heart failure**

RNA-seq from human heart samples from patients with ischemic cardiomyopathy (ICM) and controls with Non-failing hearts (NF) from GSE116250 datasets was reanalyzed, normalized, and fold-expression compared to their respective non-failing heart controls was calculated. **(a)** Volcano plots showing the Hippo signaling pathway (KEGG) significantly regulated in GSE116250 - ICM hearts (n=13) compared to NF hearts (n=14). Significantly up/down-regulated genes are represented by red dots, significant, but log2FC < 0.58 - up/down-regulated genes by blue dots, and no significant changes by gray dots (p-value < 10e-2, log2 FC > |0.58|). **b)** Bar graph representing Ingenuity Pathway Analysis (IPA) of significantly inhibited (blue bars) or activated (orange bars) signaling pathways from DEGs in GSE116250 samples. The red dotted line indicates a z-score cutoff of 2.

### Supplementary Figure 2

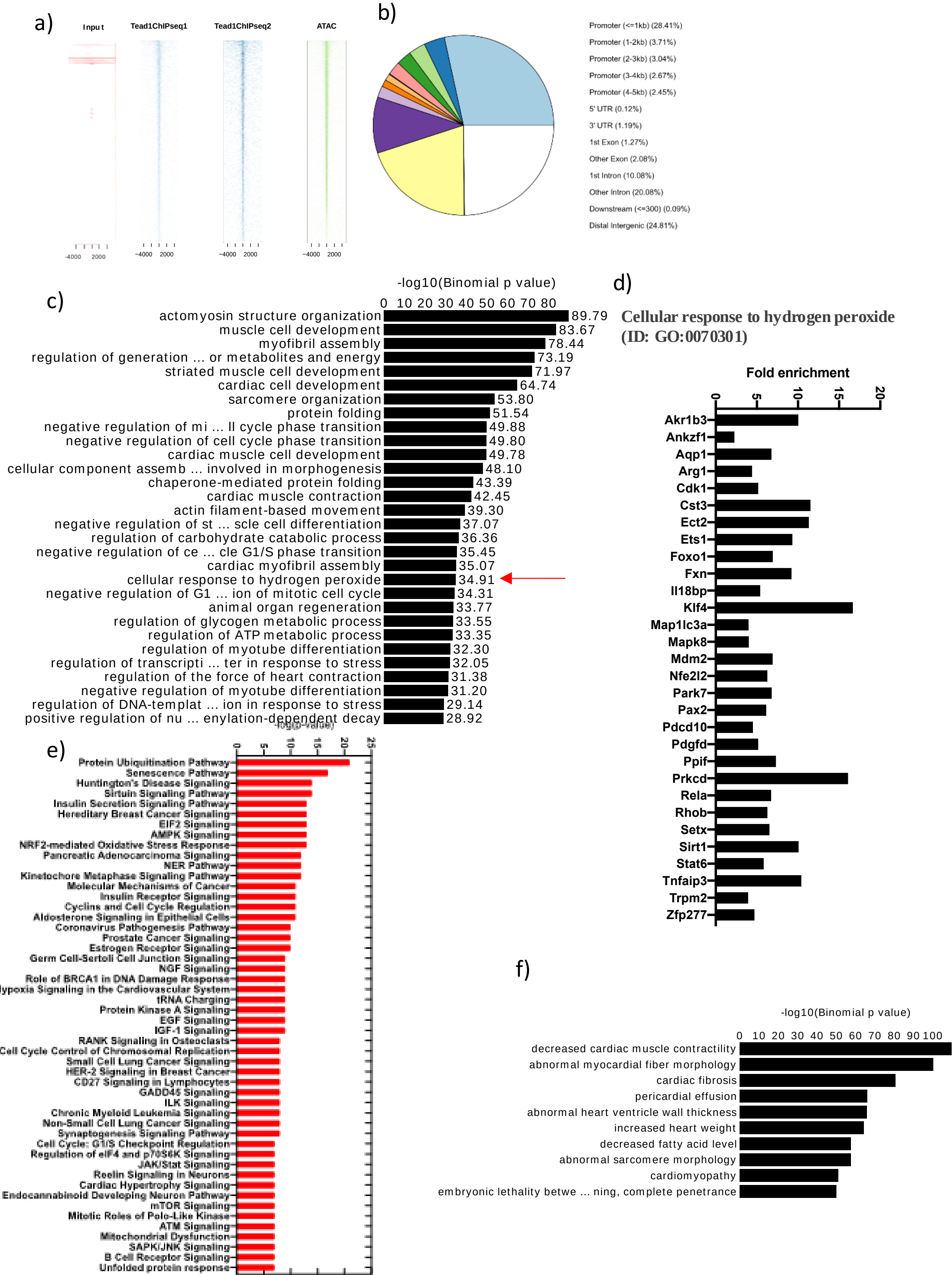

**Supplementary Figure 2: TEAD1 regulates transcription of critical cellular pathways, including anti-oxidant genes in the murine heart**. **a)** Heatmaps of normalized ChIP-seq signals for Tead1 and ATAC-seq in 10-week-old WT hearts. **b)** Genomic distribution of ATAC-seq in 10-week-old WT hearts. **c)** GREAT analysis of the top 30 significant gene ontology (GO) biological function enrichments in the Tead1 ChIP-seq, with bar heights indicating the significance of binding. **d)** Genes enriched in Tead1 ChIP-seq analysis of GO:0070301 (cellular response to hydrogen peroxide). **e)** Top 50 significantly enriched pathways identified in the ATAC-seq. **f)** GREAT analysis of the top 10 significant gene ontology (GO) mouse phenotype enrichments in the Tead1 ChIP-seq, with bar heights indicating the significance of binding.

#### Supplementary Figure 3

a)

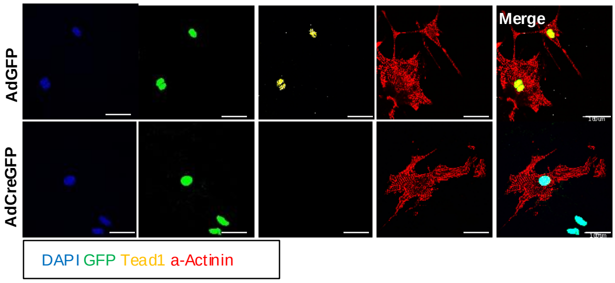

**Supplementary Figure 3: Validation of Tead1 deletion in murine neonatal cardiomyocytes. a)** Representative images of immunofluorescent staining for TEAD1 in isolated murine neonatal cardiomyocytes (iNCM) from Tead1f/f mice, 72 hours after treatment with adenoviral-GFP (AdGFP) (Top) and adenoviral-GFPCre (AdGFPCre) (Bottom).

### Supplementary Figure 4

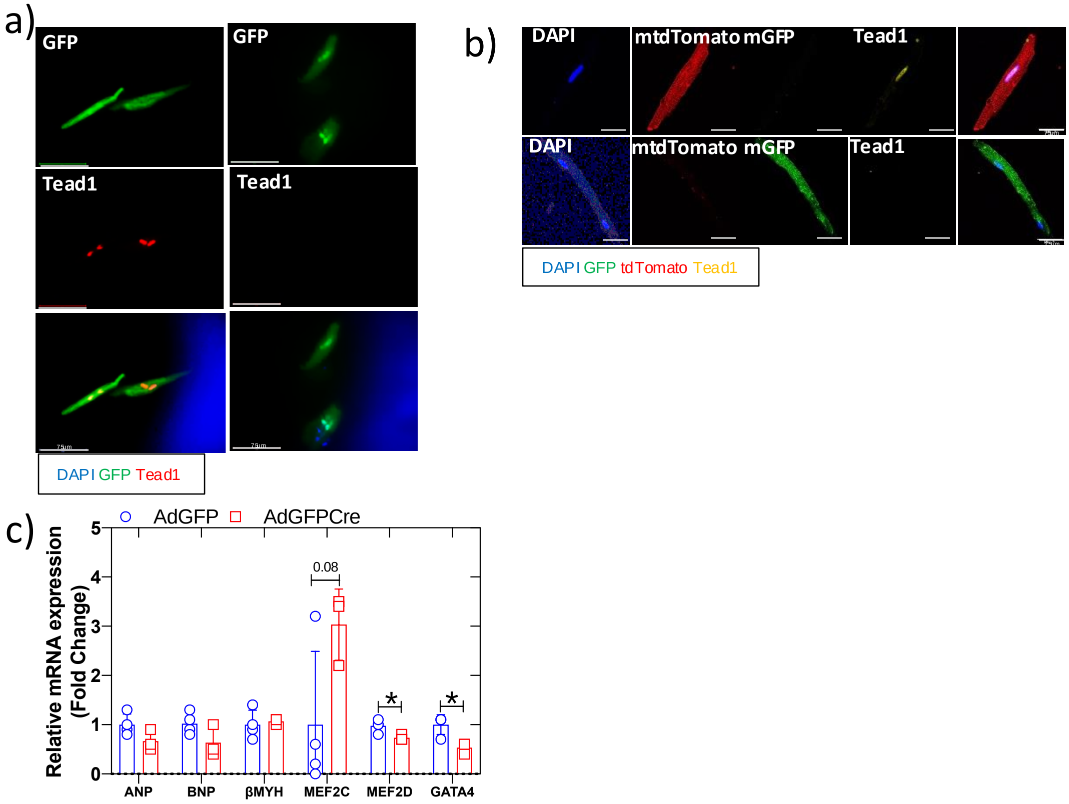

**Supplementary Figure 4: Validation of Tead1 deletion in murine adult cardiomyocytes. a)** Representative images of immunofluorescent staining for TEAD1 in isolated murine adult cardiomyocytes (iACM) from Tead1f/f mice, 72 hours after treatment with adenoviral-GFP (AdGFP) (left) and adenoviral-GFPCre (AdGFPCre) (right).

**b)** Validation of TEAD1 knockout (Tead1KO) in iACM from 10-week-old mTmG-Tead1f/f mice, after treatment with AAV9-TnTCre at postnatal day 4 (P4). Representative immunofluorescent images in these mice show tdTomato-Red positive cardiomyocytes display intact TEAD1 (yellow), while green cardiomyocytes lack TEAD1 (yellow) in the nucleus. **c)** Expression of cardiomyocyte-specific genes, including ANP, BNP, βMHC, MEF2c/d, and GATA4, performed using qRT-PCR in iACMs from Tead1f/f mice treated with adenoviral-GFP (AdGFP) and adenoviral-GFPCre (AdGFPCre) for 72 hours. Values represent fold changes compared to the control group (±SEM). Statistical significance is indicated as follows: ***$p<0.0005$; **$p<0.005$; *$p<0.05$; no symbol indicates non-significant (n = 3–6 per group). Scale bar: 75 µm.

### Supplementary Figure 5

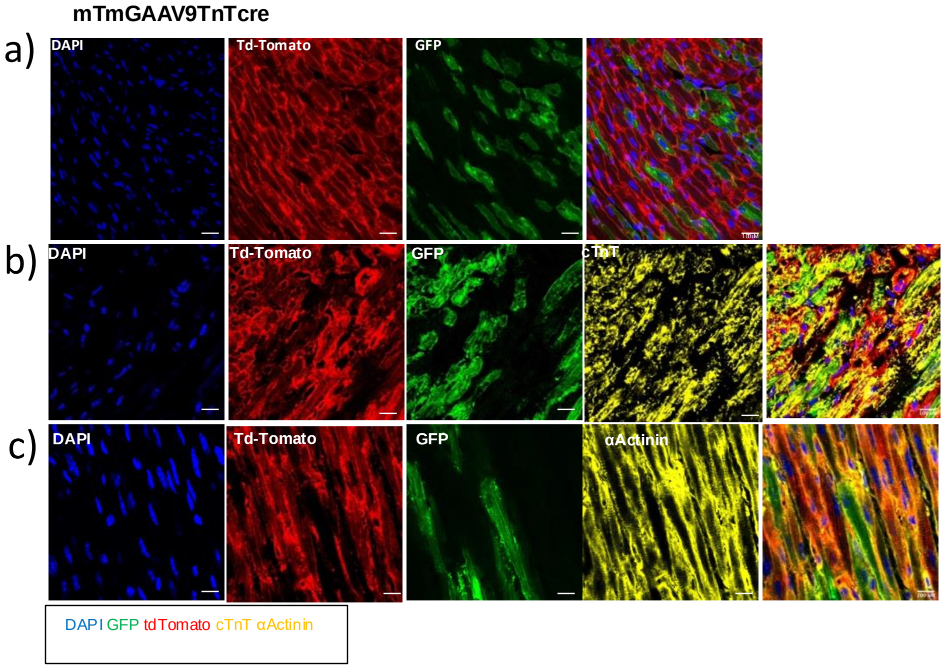

**Supplementary Figure 5: Validation of cardiomyocyte-specific Tead1 mosaic-deletion mice. a)** Harvested mouse hearts were validated for Cre activation by identifying green cells in mTmG mice treated with AAV9-TnTCre. Representative immunostaining images show that AAV9-TnTCre-injected mTmG mice have ~50% of green cardiomyocytes. **b-c)** Representative immunostaining images showing green cells with cardiomyocyte-specific markers [anti-cTroponinT (anti-cTnT) (yellow) and anti-α-actinin (sarcomeric) (anti-ACTN2) (yellow) staining]. [Nucleus-DAPI; tdTomato-mTomato; GFP-mGFP; yellow-cTnT and α-actinin immunofluorescent staining]. Scale bar: 100 µm.

### Supplementary Figure 6

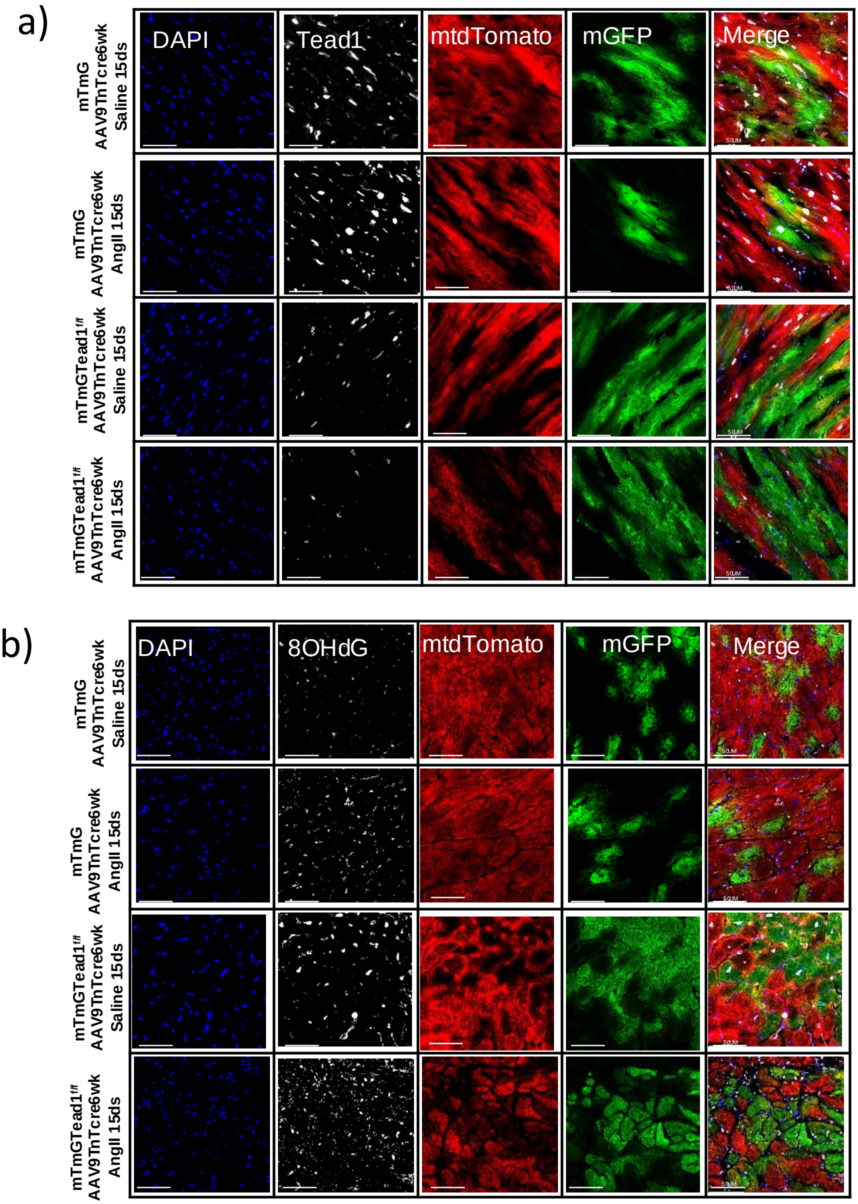

**Supplementary Figure 6: TEAD1 is required to mitigate Angiotensin II-induced oxidative stress. a)** Representative immunofluorescent images of TEAD1 mosaic hearts from mTmG-AAV9-TnTCre and mTmG-Tead1f/f-AAV9-TnTCre mice treated with Ang II or saline. [GFP-green, tdTomato-red, anti-Tead1-white, and nucleus-DAPI-blue staining]. **b)** Representative immunofluorescent images of TEAD1 mosaic hearts, showing oxidative-induced DNA damage staining by 8OHdG in mTmG-AAV9-TnTCre and mTmG-Tead1f/f-AAV9-TnTCre. [GFP-green, tdTomato-red, anti-8OHdG-white, and nucleus-DAPI-blue staining]. Scale bar: 100 µm.
